# Ancient convergence with prokaryote defense and recent adaptations to lentiviruses in primates characterize the ancestral immune factors SAMD9s

**DOI:** 10.1101/2025.05.19.654893

**Authors:** Alexandre Legrand, Rémi Demeure, Amandine Chantharath, Carine Rey, Julie Baltenneck, Cameron L.M. Gilchrist, Joana L. Rocha, Clara Loyer, Léa Picard, Andrea Cimarelli, Martin Steinegger, Francois Rousset, Peter H. Sudmant, Lucie Etienne

## Abstract

Human *SAMD9* and *SAMD9L* are duplicated genes that encode innate immune proteins restricting poxviruses and lentiviruses, such as HIV, and implicated in life-threatening genetic diseases and cancer. Here, we combined structural similarity searches, phylogenetics and population genomics with experimental assays of SAMD9/9L functions to resolve the evolutionary and functional dynamics of these immune proteins, spanning from prokaryotes to primates. We discovered structural analogs of SAMD9/9L in the anti-bacteriophage defense system Avs, resulting from convergent evolution. Further, the predicted nuclease active site was conserved in bacterial analogs and was essential for cell death functions, suggesting a fundamental role in defense across different life kingdoms. Despite this ancestral immunity, we identified genomic signatures of evolutionary arms-races in mammals, with remarkable gene copy number variations targeted by natural selection. We further unveiled that the absence of *SAMD9* in bonobos corresponds to a recent gene loss still segregating in the population. Finally, we found that chimp and bonobo SAMD9Ls have enhanced anti-HIV-1 functions, and that bonobo-specific SAMD9L polymorphisms confer increased anti-HIV-1 activity to human SAMD9L without compromising its effect on cell translation. These SAMD9/9L adaptations likely resulted from strong viral selective pressures, including by primate lentiviruses, and could contribute to lentiviral resistance in bonobos. Altogether, this study elucidates the interplay between ancient immune convergence across kingdoms and species-specific adaptations within the Avs9 and SAMD9/9L antiviral shared immunity.

**Significance statement:** The *SAMD9* gene family encodes antiviral factors of poxviruses and lentiviruses/HIV and is implicated in genetic diseases. Here, we found strong structural similarity with proteins from the Avs anti-bacteriophage system and uncovered ancient functional convergence in immune strategies between prokaryotes and metazoans. Within mammals, and more importantly in primates, we describe a highly dynamic evolutionary history of the *SAMD9* gene family that underwent adaptive episodic gene losses. Unlike humans and chimps, some bonobos lack the *SAMD9* gene entirely. Bonobos and chimps also possess unique variants of SAMD9L enhancing anti-HIV-1 activity without compromising cell functions, suggesting super-restrictors. This could also participate in shaping SIVcpz evolution and contribute to the absence of lentivirus-infected bonobos. Overall, the seeming dichotomy between the ancient evolutionary convergence in different kingdoms and recent functional adaptation within primates highlights the arms-races between key immune defense systems and viruses. This study paves the way for evolutionary medicine, where evolutionary-based discoveries may have application to human health, providing a deeper understanding of how the immune system adapts to fight viral infections over billion years of evolution.

## Introduction

The continuous arms-races between viruses and their hosts have driven the evolution of several immune defenses across all life forms ^1,2^. In humans, an emerging actor of cell-autonomous antiviral immunity is the gene family composed of the paralogs Sterile alpha motif domain-containing protein 9 and 9L (*SAMD9/9L*). These interferon-stimulated genes (ISGs) are in tandem on human chromosome 7 and encode large, multi-domain proteins with antiviral properties against poxviruses and lentiviruses ^3–5^. SAMD9 and SAMD9L inhibit cellular and viral protein translation ^6–8^, acting through an essential ribonuclease site in the AlbA_2 domain ^5,9^. They are also modulators of endosomal trafficking ^10,11^ and SAMD9 was recently identified as a nucleic acid sensor ^12^. Deleterious germline mutations in human SAMD9 or SAMD9L dysregulate their activity leading to severe life-threatening genetic syndromes, such as MIRAGE (myelodysplasia, infection, restriction of growth, adrenal hypoplasia, genital phenotypes, and enteropathy) ^11^, SAMD9L-associated autoinflammatory disease (SAAD) ^13^, ataxia-pancytopenia (ATXPC) ^14^ and normophosphatemic familial tumoral calcinosis (NFTC) ^15^. Although the *SAMD9* gene family extends beyond humans, its evolutionary history and functional diversification remain largely unexplored.

In the past few years, some human immune genes were shown to have a deep evolutionary origin in bacterial defenses against phages ^16^. For example, major eukaryotic antiviral immune sensors or effectors, such as cGAS or Viperin, have originated from prokaryotic antiviral systems ^16–19^. Ancestral immunity therefore broadly defines shared immune defenses between prokaryotes and eukaryotes through the presence of conserved immune modules (domains or proteins) ^18^, that results from horizontal gene transfer, convergent evolution or vertical inheritance.

SAMD9 and SAMD9L are large multidomain proteins, which are presumed members of the STAND (signal transduction ATPases with numerous domains) superfamily ^20^. SAMD9s are composed of (from N-to C-terminal): a Sterile Alpha Motif (SAM), a Schlafen (SLFN)-like AlbA_2 domain with nuclease site, a predicted SIR2, a predicted P-loop NTPase, predicted tetratricopeptide repeats (TPRs) and an oligonucleotide/oligosaccharide-binding OB-fold. All are predicted domains, except the AlbA_2 which structure was recently solved ^21^. Computational analyses suggested the presence of homologs of SAMD9 domains in other animals and bacteria ^20^. However, the ancient evolutionary and functional history of SAMD9s and its potential link to immunity outside mammals remain unknown.

In mammals, the gene family consists of two paralogs, *SAMD9* and *SAMD9L*, which originally duplicated in placentals and have evolved under positive selection ^22^, suggesting past molecular arms-races with pathogens. Interestingly, these mammalian paralogs exhibit both functional redundancy and divergence in their antiviral functions. Both are restriction factors against poxviruses, but with species-specific variations in their susceptibility to poxviral countermeasures ^4^. In humans, they display different functions in human immunodeficiency virus (HIV) and lentiviral infections: SAMD9L is antiviral, while SAMD9 has no, or a proviral, effect ^5^. Lentiviruses naturally infect most African non-human primate species (SIVs, simian immunodeficiency viruses), with the notable exception of some species such as bonobos ^23–26^. SIVcpz from chimpanzees is at the origin of the HIV-1 group M, responsible for the AIDS pandemic ^27^. Lentiviruses have coevolved with primates for million years and have been selective drivers of adaptation in primate antiviral defense factors, such as APOBEC3G, Tetherin/BST-2 or TRIM5 ^28–30^. These species- and lineage-specific adaptations have further shaped host molecular barriers to cross-species transmissions ^31,32^.

In this study, we combined AI-predicted structural similarity searches, phylogenetics and population genomics with experimental assays of SAMD9/9L functions in prokaryotes and great apes to resolve the evolutionary and functional dynamics of this antiviral system across scales. We notably found that SAMD9s and some prokaryotic Antiviral STAND (Avs) originated from convergent evolution and depend on AlbA_2 domain for their activity, and that SAMD9s have evolved under recurrent diversifying evolution in mammals by genomic structural variation, notably gene losses and adaptive polymorphisms to lentiviruses in great apes.

## Results

### SAMD9 bacterial structural analogs with conserved multidomain architecture and predicted antiphage activity

Benefiting from recent advances in structural similarity methods, we investigated the evolutionary history of *SAMD9s* across the tree of life. We first used Foldseek ^33^, which allows fast protein structure alignment and search, with the AlphaFold-predicted SAMD9/9L structures and amino acid sequences as inputs (details in Methods). We identified 238 hits that share high structural similarity, mainly belonging to bacteria and metazoa (30% Template Modeling TM-score and 80% coverage cut-off). Analysis of bacterial hits with DefenseFinder ^34^, which performs Hidden Markov Model (HMM) searches against a database of prokaryotic antiviral systems, showed that some of these hits belong to the Avs family (antiviral STANDs) of anti-phage immune proteins ^35–38^. Avs proteins are NLR-like proteins that sense viral phage infections by their C-terminal sensor domains and perform specific antiviral functions through their N-terminal effector domains. The effector domain may have diverse specific activities, such as ATP degradation via a PNP domain, or NAD+ depletion via a SIR2 domain ^17,37,39,40^. Here, we found that SAMD9 shares strong structural similarity with Avs5 from *Desulfobacula sp* and from a newly identified protein, which we named “Avs6”, from *Labedella endophytica* (Uniprot IDs: A0A1F9N8W4 and A0A3S0VH60, respectively) (Fig. 1A). However, Avs5 and Avs6 lack the N-terminal AlbA_2 domain from SAMD9, with Avs6 instead encoding a PNP domain predicted to degrade ATP molecules (Fig. 1A). Remarkably, our analyses also identified other bacterial proteins sharing strong structural similarity with human SAMD9/9L on up to 88% of the protein length (e.g. 1407/1589 aa total in SAMD9 and 1400/1584 in SAMD9L for A0A7T4VS34 from *Pseudomonas fluorescens*). This similarity covers all its functional domains, except the first N-terminal SAM, which seems exclusive to eukaryotes ^41^. Despite the low amino acid sequence identity (around 15%), the structural similarity is highly significant (TM-score up to 0.56; Fig. S1A) ^42^. We therefore propose “Avs9” as a name for bacterial Avs proteins harboring the same domain composition as SAMD9/9L at the exception of the SAM domain, referring to the latter gene family name (Fig. 1A). Furthermore, our analysis identified hundreds of uncharacterized proteins across various domains of life: metazoa, bacteria and archaea. It is however notable that, although STAND proteins are also present in plantae and fungi ^43,44^, we did not retrieve any significant hits in these kingdoms of life, neither in algae nor in protozoa. Further searches with various inputs, including Avs9, and more relaxed parameters did not recover any.

**Figure 1.**
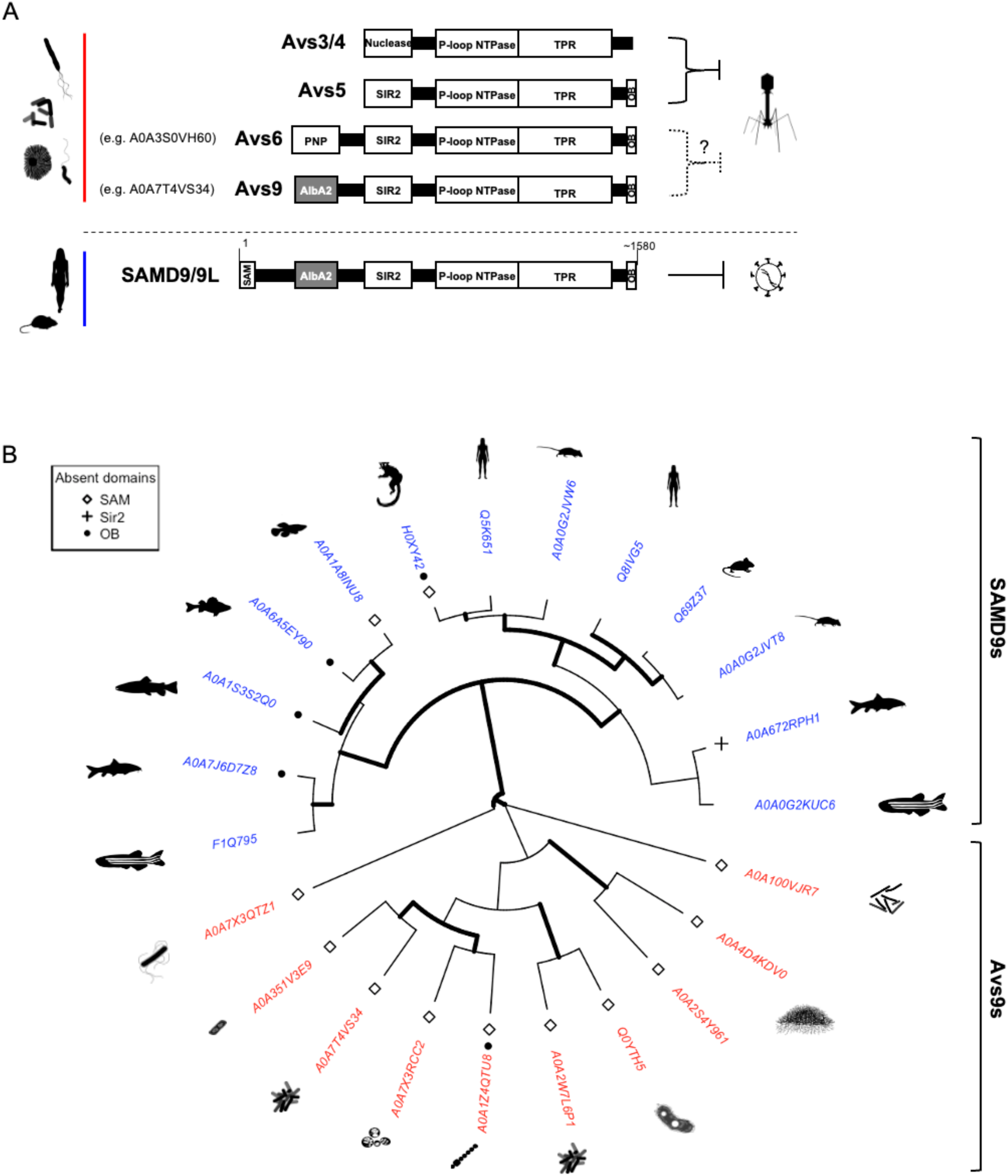
Structural homology analyses show strong similarity between SAMD9 gene family and Avs antiphage proteins. **A,** Structural homology searches of human SAMD9/9L in prokaryotes identify known and predicted Avs antiviral systems. Linear representation of the multi-domain organization of proteins with strong SAMD9/9L structural similarity showing a common conserved organization, on more than 1,300 aa for Avs9. Avs3, Avs4 and Avs5 were previously identified ^35,37^, while Avs6 and Avs9 are proposed names of newly identified Avs members (in parentheses, representative members with protein UniProt ID). **B,** Circular ultrametric tree representing structural clustering of SAMD9s and Avs from FoldMason multiple structure alignment showing widespread and diverse SAMD9/9L structural analogs in bacteria (red) or metazoan (blue) SAMD9s. Tree rooting is only for representation purposes. Statistical supports are from 1000 bootstrap replicates (values above 90 are represented by thick lines). Protein with an absent domain have either a square, a cross or a filled circle at the tip of the corresponding branch, for the absence of SAM, SIR2 or OB, respectively. Species silhouettes are from https://www.phylopic.org.

To investigate the evolutionary history of the structural analogs, we performed maximum likelihood phylogenetic analyses, using IQ-TREE, of both the sequence and the structural alignments of these proteins (from Muscle and FoldMason, respectively) (Fig. S1B). We found that SAMD9s and Avs9s were in two distinct clades falling within the same lineage. However, because the proteins bear different domain compositions, we next performed analyses for specific domains, individually (Fig. S1C). Starting with the central P-loop NTPase domain, which is a domain common to all hits, we similarly found that Avs9 and SAMD9 clades fell into the same lineage in the phylogeny (Fig. S1D). Next, we only kept proteins encoding for an AlbA_2 effector domain, resulting in 23 hits (i.e. the small number of hits are due to the AlphaFold Database clustered at 50% identity): 13 eukaryotic SAMD9/9Ls and 10 bacterial Avs9s. The resulting tree topology also showed two clades corresponding to SAMD9s and Avs9s (Fig. 1B). We further observed scattered absence of SAM, OB-fold or SIR2 domains (Fig. 1B). While this suggests that these proteins have a propensity for domain modularity and that these domain losses could be tolerated, the functional impact of such modulation would require further investigations. Overall, the extensive structural similarity of SAMD9- and Avs9-like proteins observed across such a vast evolutionary range is remarkable. It suggests a strong selective advantage driving the structural and functional conservation of this putative immune antiviral system from bacteria to mammals.

### Eukaryotic SAMD9s and bacterial Avs9s result from convergent evolution

To determine whether eukaryotic SAMD9s result from the ancestral acquisition of a full-length bacterial analog (such as Avs9) or emerged through convergent evolution, we performed new structural similarity searches for AlbA_2-containing proteins and studied the phylogenetic distribution of this effector domain across the tree of life (Fig. 2A-B). First, we found that AlbA_2 was not only present in Avs9-like prokaryotic proteins, but was widely represented in bacteria. Concerning eukaryotes, well-known AlbA_2-containing proteins included members of SAMD9 and SLFN antiviral immune factors, which, in our analysis, fell in distinct clades (except for SLFNL1, which is distantly related to other SLFNs ^45^ and whose evolutionary history was uncertain), suggesting different evolutionary origins (Fig. 2B, S2B-C). As for the ten identified Avs9s, they all clustered within a single clade that was distant from eukaryotic SAMD9s (Fig. 2B, S2B-C), showing that full-length Avs9- and SAMD9-like proteins did not originate from a single common ancestor. Overall, the presence of evolutionarily distant AlbA_2 domains together with a strong structural similarity and a similar domain organization support a model in which bacterial Avs9s and eukaryotic SAMD9s evolved similar domain architectures through convergent evolution driven by analogous selective pressures.

**Figure 2.**
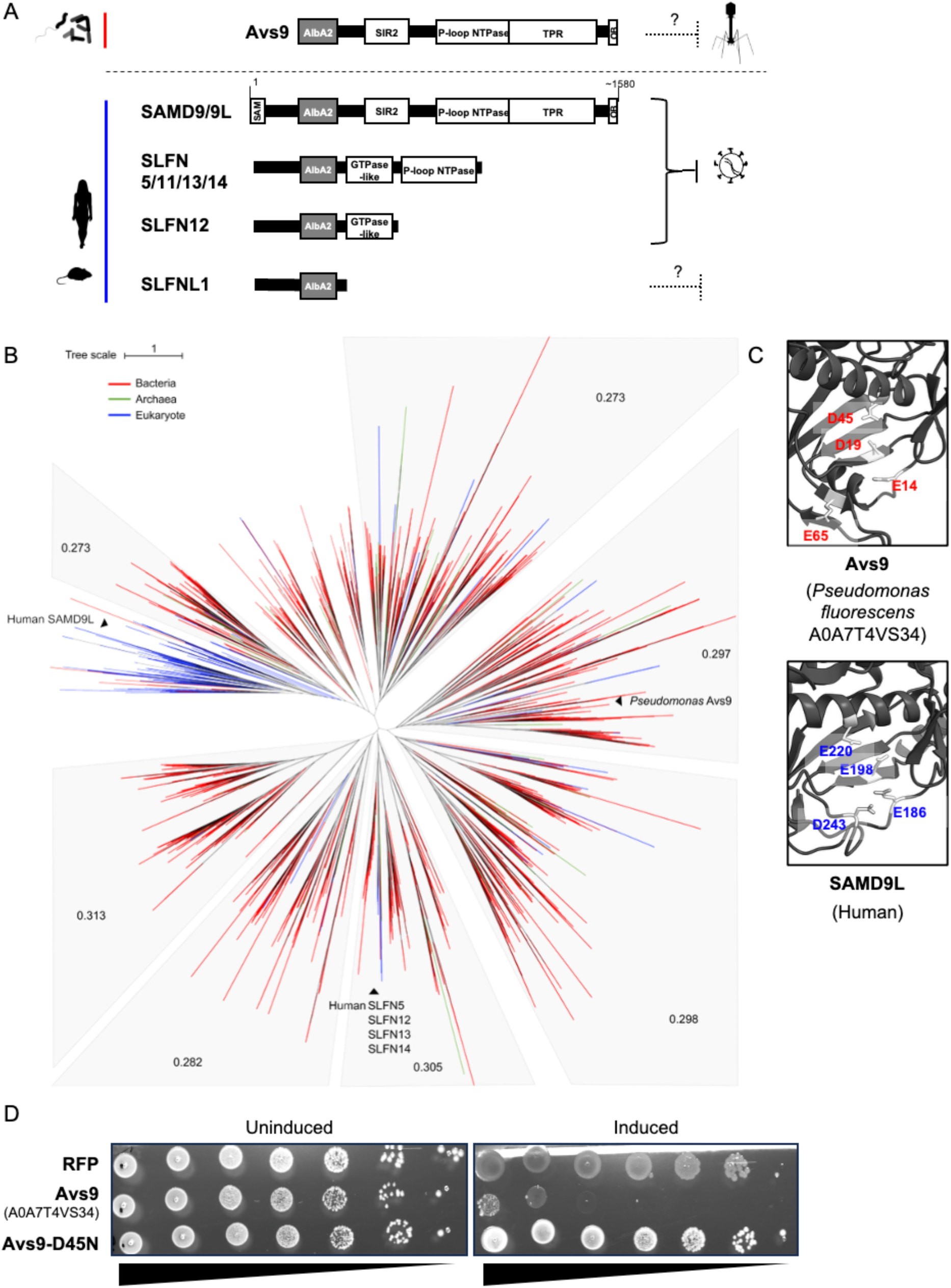
Avs9 is part of prokaryotic defense systems and induces cell death in bacteria, through its SAMD9/SLFN-analogous active site in the AlbA_2 domain. **A,** Linear representation of the multi-domain organization of key proteins bearing an AlbA_2 domain: prokaryotic Avs9 and proteins from the SAMD9s and SLFNs**. B,** Unrooted phylogenetic tree of a multiple sequence alignment of AlbA_2 domains detected in proteins from kingdoms of life, visualized on iTOL. Shown for each clade is the defense score (i.e. the fraction of bacterial AlbA_2 domains encoded in the vicinity of known defense systems; see Methods). **C**, Predicted structures of AlbA_2 domains of *Pseudomonas fluorescens* A0A7T4VS34 (Avs9) and human SAMD9L, showing conserved SLFN-like nuclease catalytic site. Residues forming the catalytic sites are in clear grey with their coordinates in red or blue. **D,** Ten-fold serial dilutions of *E. coli* cells transformed with plasmids encoding either Avs9, Avs9-D45N or RFP as a control, with induction (100µM IPTG) or without induction (1% glucose). Shown are photos of bacterial drops.

Antiphage defense systems tend to physically cluster in defense islands of bacterial genomes ^46^. To investigate the possible immune function of bacterial AlbA_2 domains, we used DefenseFinder to evaluate their propensity to be encoded in genomic neighborhoods of known defense systems. We found a significant association of bacterial AlbA_2 domains with defense systems in all tested clades, including Avs9 (Fig. 2B), strongly suggesting that their primary function is immune defense.

### AlbA_2 domain is a shared effector determinant of SAMD9s’ and Avs9s’ activity

To investigate the function of bacterial Avs9 proteins, we first looked for the conservation of the AlbA_2 catalytic site. This active ribonuclease site is composed of three to four negatively charged residues, which are necessary for SAMD9/9L and SLFN11/12/13/14 antiviral activities ^5,9,47–52^. Here, we found similar residues and structure in Avs9 AlbA_2 (Fig. 2C: E14, D19, D45, E65, Fig. S2A), suggesting the presence of similar catalytically active site and function.

Second, to investigate Avs9 activity *in vivo*, we selected and synthesized an Avs9 from *Pseudomonas fluorescens* (A0A7T4VS34), and we cloned its full-length gene, or the AlbA_2 domain alone, under an inducible promoter into *E. coli*. Expression of full-length Avs9 (Fig. 2D), or the AlbA_2 domain alone (Fig. S2D), led to major cell death upon mild induction at 25°C. Interestingly, the *Pseudomonas fluorescens* Avs9 cell death induction was abolished by introducing a single-residue mutation in the AlbA_2 predicted catalytic site (D45N; Fig. 2D, S2D). Of note, an equivalent mutation in human SAMD9/9L and SLFN AlbA_2 also abolishes their activity. Therefore, our results strongly suggest a cell-killing defense function of bacterial Avs9, dependent on AlbA_2 and its predicted nuclease active site.

### Ancient duplication followed by frequent copy number variations (CNVs) of the *SAMD9* gene family in mammals

Although *SAMD9s* are present in multiple vertebrate hosts (Fig. 1B), the *SAMD9* duplication at the origin of the human paralogous *SAMD9* and *SAMD9L* genes seems more recent. This duplication was previously described in placental Eutherians, after the divergence of Marsupials from Placentals ^22^. To address the evolutionary history of the duplicated *SAMD9/9L*, we first performed genomic analyses of the *SAMD9* gene family locus in various vertebrate species focusing on mammals: 38 ungulates, 22 chiropterans, 45 carnivores, 34 primates, 41 rodents, and 27 additional species from other mammalian and non-mammalian vertebrate orders (Fig. 3A). We found that all these analyzed vertebrate species, except monotremes (n=2), exhibited at least one gene from the *SAMD9* gene family (Fig. 3A).

**Figure 3.**
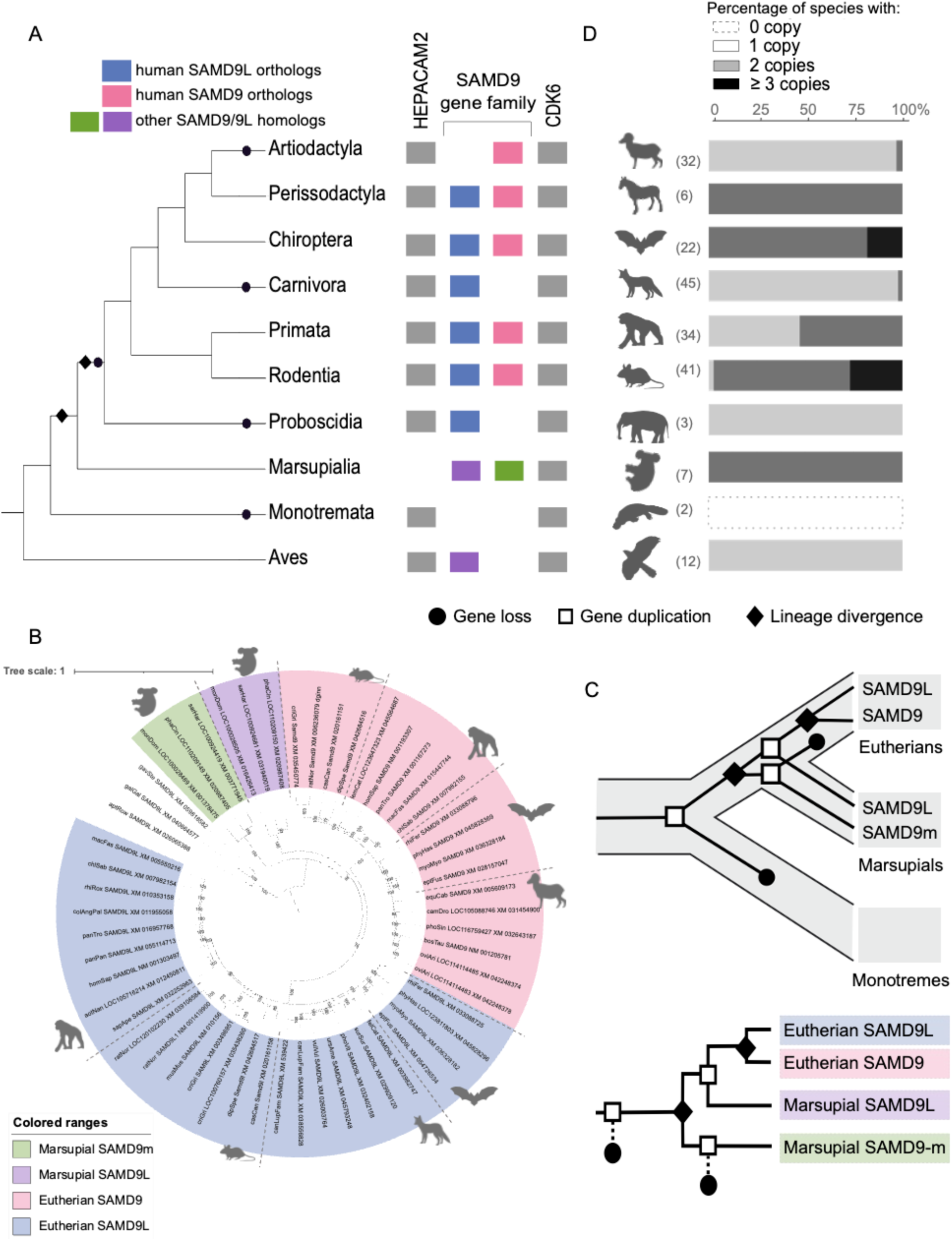
Ancient duplication followed by frequent copy number variations of the *SAMD9* gene family in mammals. **A,** Representation of the *SAMD9* gene family locus in each mammalian order and other vertebrates. Order cladogram is presented on the left with diamonds and rounds on the branches representing events of gene gain and loss, respectively, in the major lineages only. Colored rectangles represent the gene members of the *SAMD9* gene family with orthologs to human *SAMD9L* and *SAMD9* in blue and pink, respectively. Grey rectangles represent adjacent syntenic genes. **B,** Maximum likelihood phylogenetic tree (IQ-TREE, GTR+F+I+G4 substitution model) generated with selected SAMD9/9L homologs from Eutherians, Marsupials, as well as Aves and Amphibians, used as outgroups. Bootstraps are from 1,000 replicates. The complete tree is shown in Fig. S3A. The scale bar represents the number of substitutions per site. **C,** Schematic diagrams of the origin of *SAMD9/9L* duplication in mammals. Top, grey tree in the background represents the species evolution of the three mammalian groups. Black tree inside the grey one represents the evolution of the *SAMD9/9L* gene tree. Bottom, Alternative representation with the gene tree cladogram. Legend is embedded. **D,** For each order, copy number variations (CNVs) in the *SAMD9/9L* gene family are indicated by histograms. Alignments and trees are available (see Data Availability). Bar colors (grey scale) represent the number of *SAMD9/9L* copies (legend embedded). Bar lengths indicate the proportion of genomes with the indicated number of copies for each order, with the total number of analyzed genomes presented in parentheses for each order (aligned to panel A). Species silhouettes are from https://www.phylopic.org.

Furthermore, thanks to genomic sequence advances and contrary to prior findings^22^, our observations indicated that Marsupials also possess two copies and therefore the gene duplication of *SAMD9s* was not restricted to Eutherians (Fig 3A-B). To elucidate the origin of *SAMD9/9L* in mammals, we performed phylogenetic analyses from 301 homolog sequences of 189 mammalian species spanning 160 million years of divergence (using NCBI blastn implemented in DGINN ^53^) and from 18 non-mammalian vertebrate species (i.e. 12 aves and 6 amphibians) as outgroups (Fig. 3B with selected species and IQ-TREE tree, Fig. S3A-B for the complete IQ-TREE and PhyML trees using sequences from Dataset S1; of note: PhyML trees were similar to IQ-TREE trees at key branches). Remarkably, despite the presence of two copies in marsupial genomes (Fig. 3A), one of them, named *SAMD9m*, did not group with the Eutherian *SAMD9* or *SAMD9L* clade in the homologous gene tree, but branched outside (Fig. 3B, statistically significance assessed from 1,000 bootstrap replicates). Therefore, it is likely that the marsupial *SAMD9m* resulted from an ancestral independent duplication predating the divergence of Marsupials and Eutherians (Fig. 3B). Following this divergence, one copy was potentially lost in Eutherians, followed by a subsequent duplication event (Fig. 3C). Alternatively, other hypotheses may involve a single duplication event followed by gene conversion within the two Eutherian copies, or loss of a copy through incomplete lineage sorting (ILS).

In placentals, following the duplication event that gave rise to *SAMD9* and *SAMD9L* orthologs, we identified at least 10 independent losses of one of the two paralogs throughout mammalian evolution (5 losses in the main represented lineages of Fig. 3A, as well as additional losses within orders: Fig. 3D, S3C). Notably, artiodactyls experienced the loss of *SAMD9L*, while carnivores lost *SAMD9*. Furthermore, although the synteny and the copy numbers remained largely conserved within the divergence of the latter two groups, radiations of primates, bats and rodents, exhibited many SAMD9/9L gene losses and some duplication events (Fig. 3D, Fig. S3C, see Data availability).

### Ancient and recent unfixed gene losses, as well as lineage-specific positive selection, during primate SAMD9/9L evolution

Genomic structural changes, such as gene duplication and loss (i.e. CNVs), alongside mutation and recombination events, are an important source of genetic diversity upon which natural selection can act, and thus have strong adaptive potential during virus-host arms races ^54,55^. Such genomic adaptations during mammalian evolution have been rampant in response to past lentiviral or poxviral epidemics ^30,56^. Because of (i) the very high rate of CNVs in primate *SAMD9/9Ls*, (ii) the co-evolution of primates with lentiviruses for millions of years ^32^, and (iii) the potential lentivirus-driven adaptation of SAMD9/9L ^5^, we performed in-depth phylogenomic, genetic and positive selection analyses in primates.

We identified at least four independent gene loss events during primate evolution (Fig. 4A). *SAMD9L* was lost in prosimians, while bonobos and the common ancestors of Platyrrhini and *Colobinae* experienced the independent loss of *SAMD9*. Genomic analyses showed that *SAMD9* loss in Platyrrhini resulted from the complete loss of the SAMD9 genomic locus in the common ancestor. In contrast, *SAMD9* loss in *Colobinae* occurred through a different genetic mechanism by single nucleotide changes introducing multiple premature stop codons in the coding sequence (Fig. 4A, S4A).

**Figure 4.**
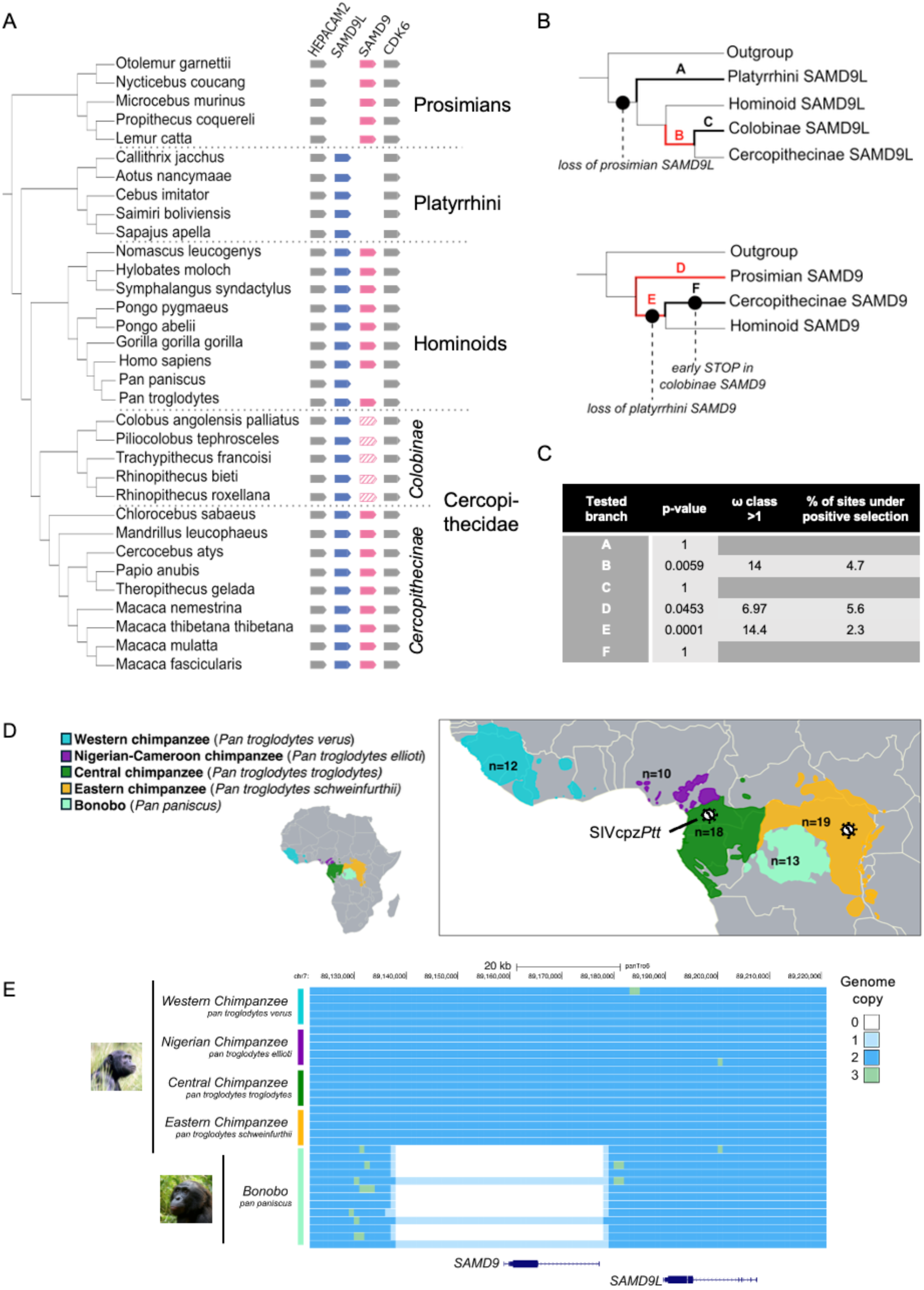
Ancient and recent unfixed gene losses in primates with lineage-specific adaptation. **A,** Representation of the *SAMD9* genomic locus from primate genomes. Species cladogram is pre-sented on the left. *SAMD9L* and *SAMD9* are in blue and pink, respectively. Adjacent syntenic genes are in grey. Genes containing early stop codons are hatched. **B,** Simplified primate gene cladograms of *SAMD9L* (top) and *SAMD9* (bottom) showing, in bold, the branches tested for positive selection with aBSREL (HYPHY) and, in red, the branches under significant positive selection (p-value<0.05). The “outgroup” branch corresponds to *Cricetulus griseus* (criGri) and *Tupaia chinensis* (tupChi) and is shown here as a rooted tree for representation purposes. **C,** For each branch tested, the table recapit-ulates the statistical significance of positive selection assessed by LRTs with the Holm-Bonferroni cor-rection, the estimated dN/dS value of the sites in the class under positive selection, and the proportion of sites under positive selection (%). **D,** Africa map with inset representing the current geographic ranges of *Pan* populations and their status regarding natural SIV infections. Numbers of individuals studied for their whole-genome sequences are indicated for each (sub)species. Populations naturally infected by SIVcpz are highlighted by a virus symbol. **E,** *SAMD9/9L* genomic locus alignment among *Pan* individuals showing recent unfixed loss of *SAMD9* in the bonobo population.

We next specifically tested whether episodes of positive selection occurred in SAMD9 or SAMD9L in primate lineages that experienced paralog loss. We performed targeted branch-specific analyses, using the adaptive Branch-Site Random Effects Likelihood (aBSREL), testing specifically lineages associated with gene loss or paralog retention (“tested branch”, also known as “foreground” branches) in the SAMD9 or the SAMD9L gene trees (others branches were set as “background branches”) ^57^(Fig. 4B-C). Our analysis suggested that several primate lineages that lost either *SAMD9* or *SAMD9L* were the targets of episodic positive selection (Fig. 4B-C). The observed gene losses within primates, coupled with evidence of episodic positive selection, either prior to the copy loss or in the remaining copy, suggest ancient adaptive response to pathogen challenges. These losses could represent evolutionary trade-offs, where the benefits of losing a gene (potentially escaping viral antagonism or hijacking) outweighed the costs.

Bonobos and chimpanzees are human’s closest living relatives with a genetic divergence of approximatively 1.3% with humans, and only of 0.4% amongst themselves ^58^. Despite this strong proximity, bonobos possess a unique genetic distinction, with a 41.46 kb deletion in the *SAMD9/9L* locus. In fact, they stand out as the sole hominid with a single copy of the *SAMD9* gene family, retaining only *SAMD9L* ^59^. To characterize the recent loss of *SAMD9*, we investigated the prevalence of the 41.46 kb deletion in the *SAMD9/9L* locus in bonobos at a population level (*Pan paniscus*; n=13) in a joint analysis that additionally included all currently recognized chimpanzee subspecies or populations (*Pan troglodytes spp*; n= 59) (Fig. 4D). We found that, among 13 individuals, 10 bonobos exhibited the same deletion with the complete absence of the *SAMD9* gene, indicating common homozygous genomic deletion (Fig. 4E). However, the remainder three bonobos presented this deletion on a single chromosome, representing a heterozygous genomic absence of *SAMD9* (Fig. 4E). Importantly, the analyzed bonobo individuals are not related ^60^, suggesting that, despite the small sample size, the heterogeneous genomic makeup in SAMD9/9L may be representative of the population. This was in sharp contrast with chimpanzees, which had *SAMD9^+/+^* present in all 59 individuals, a pattern shared with humans (Fig. 4E). This suggests a recent *SAMD9* genomic loss event specific to the bonobo lineage still segregating in the population, potentially impacting bonobo’s immunity.

### Chimp and bonobo (*Pan*) SAMD9Ls have an increased anti-HIV-1 activity compared to human SAMD9L

Beyond the genomic loss that occurred in the *SAMD9/9L* locus during hominid evolution, we investigated the evolutionary divergence of *SAMD9* and *SAMD9L* at the genetic level. We therefore analyzed the non-synonymous single nucleotide polymorphisms (SNPs) within the coding sequences of both genes for the 72 *Pan* individuals, using panTro6 genome as reference, as well as for more than 4,000 humans (Fig. 5A, Fig. S5A-B). For SAMD9, we found very few non-synonymous SNPs amongst *Pan*. This amino-acid conservation was particularly exemplified in the 12 Eastern chimps (*P. t. verus*) and in the 3 bonobos encoding a single SAMD9 copy, as they encoded an identical protein sequence (Fig. 5A). In SAMD9L, we found a more widespread distribution of missense SNPs in chimps and bonobos (Fig. 5A). Remarkably, we found two specific variants in SAMD9L that stood out due to their high frequency in the bonobo population. All bonobos (n=13) encoded for a homozygous Serine (S) at position 90, and 9 out of 13 bonobos encoded an Arginine (R) at position 1446 (either homozygous or heterozygous) (Fig. 5A, S5A). Of note, there were no SAMD9L SNPs differentiating SAMD9^+/-^ and SAMD9^-/-^ bonobos. Intriguingly, chimps and all other primates analyzed to date, including the 4,099 human genomes, encoded a Leucine L90 and a Lysine K1446 (Fig. 5A, Fig S5A-C), suggesting that these variants are specific to bonobos amongst primates. Furthermore, compared to bonobos, Eastern and Central chimps showed a more widespread distribution of SNPs within SAMD9L, and no missense polymorphisms appeared at high frequency in the chimps (Fig. 5A, S5A).

**Figure 5.**
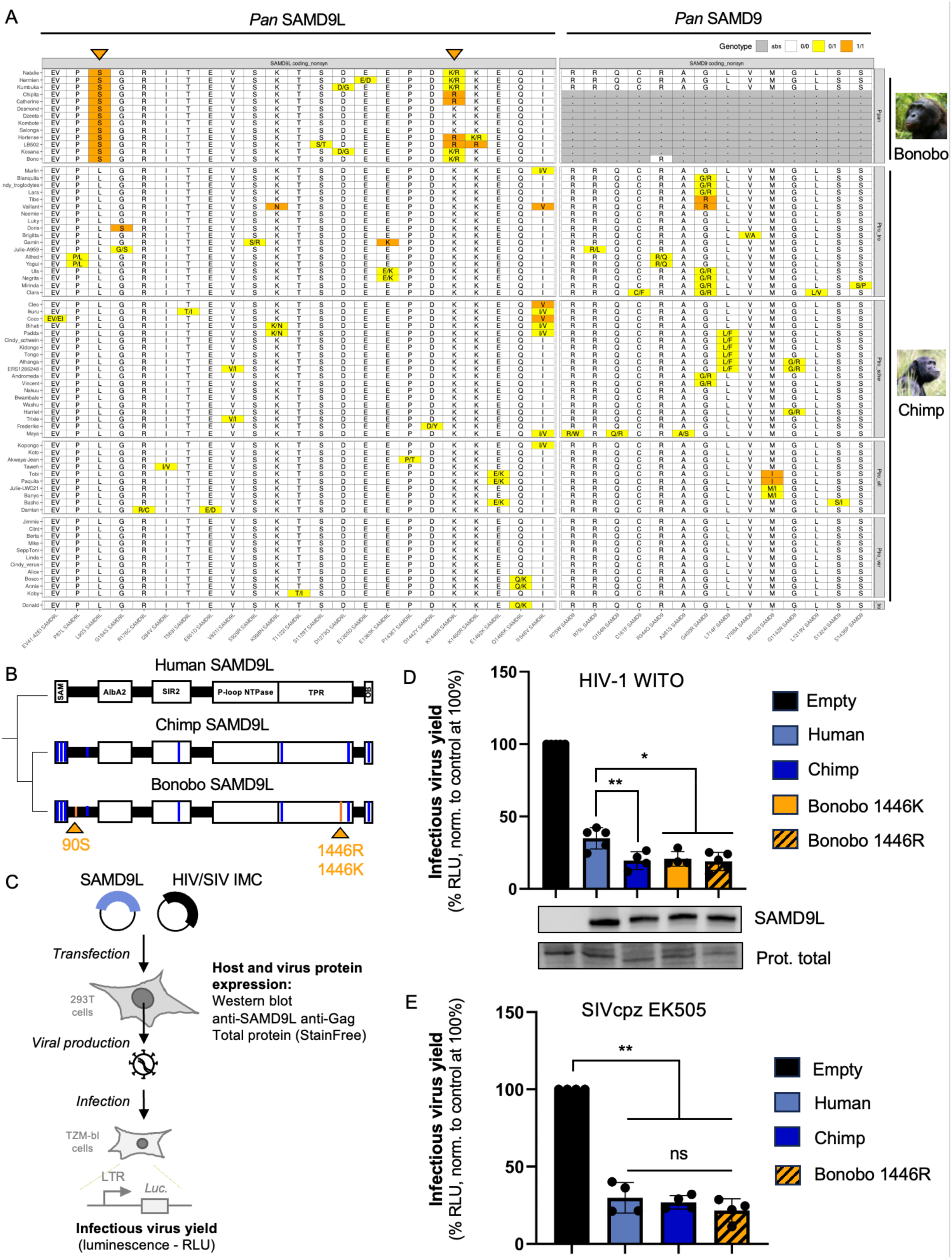
Chimp and bonobo (*Pan*) SAMD9L major variants have an increased restrictive activity against HIV-1 pWITO, but not SIVcpz EK505, compared to human SAMD9L. **A,** Polymorphisms impacting SAMD9 and SAMD9L amino acid coding sequences among the *Pan* populations, with chimp PanTro6 as reference. Highly frequent bonobo-specific SNPs in SAMD9L are highlighted by orange triangles at the top. See Fig. S5 for details, human polymorphisms, and comparative analyses with other primate species. **B,** Location of the chimp and bonobo specific variants as compared to human SAMD9L on the 2D predicted protein domain structure (n=8 for chimps versus humans, n=9-10 for bonobos versus humans). **C,** Experimental set-up to investigate chimp and bonobo SAMD9L restriction on replication of full-length HIV-1 and SIVcpz infectious molecular clones (IMC). **D,** Relative infectious virus yields of HIV-1 pWITO in indicated SAMD9L conditions, normalized to the empty control (Experimental setup in C). The two most frequent variants at position 1446 in the bonobo populations were functionally tested. Results from five independent biological replicates. Statistics were performed using the ratio paired t-test versus the human SAMD9L condition (**, p value <0.005; *, p value < 0.05). Of note, all SAMD9Ls significantly restricted HIV-1 pWITO as compared to the empty control (p<0.005). Below, Western-blot analysis in the producer cells showing similar protein expression of SAMD9Ls. Loading control is from total protein (Prestained gels). **E,** Similar experiment, as in C-D, with SIVcpz EK505 strain. ns, not significant.

Human SAMD9L inhibits cellular protein synthesis and restricts lentiviral HIV-1 infection ^5,8,61,62^. This is particularly interesting in the context of natural lentiviral infections in hominids, where humans and two chimp subspecies are infected by HIV-1 and SIVcpz, respectively. Yet, two chimp subspecies (Eastern and Nigerian-Cameroon chimps) and bonobos have no evidence of modern natural lentiviral infections ^25–27,63^ (Fig. 4D). Furthermore, pandemic HIV-1 in humans originated from cross-species transmission of SIVcpzPtt from Central chimps (*Pan troglodytes troglodytes*) ^27,64^.

We therefore determined the functional consequences on lentiviral infections of the genotypic differences between human, chimp and bonobo SAMD9Ls. We cloned the native chimp SAMD9L and the two major variants of bonobo SAMD9L (bonobo R1446 and bonobo K1446) in an expression plasmid to compare them with human SAMD9L (Fig. 5B-C). Of note, the chimp SAMD9L has 8 amino acid (aa) changes compared to the human one (Fig. 5B). We investigated their functions in the context of HIV-1 replication, as in^5^, as well as SIVcpzPtt replication (SIVcpz EK505, a kind gift from BH Hahn ^27,65^). Briefly, we co-transfected 293T cells with the mCherry-SAMD9L plasmids along with an infectious molecular clone (IMC) encoding a transmitted/founder HIV-1 natural strain (pWITO, as in^5^) or an SIVcpzPtt strain (EK505). Two days later, we measured viral and cell protein expression in the producer cells (Fig. 5C-E, S5D) and the infectious virus yield by Tzm-bl reporter assay (Fig. 5C-E, S5D). Remarkably, although all ectopic SAMD9Ls were expressed at similar levels, we found that chimp and bonobo SAMD9Ls had a significant increase in anti-HIV-1 pWITO activity compared to human SAMD9L (Fig. 5E-F, p<0.005), suggesting some species-specificity.

By testing effects on SIVcpzPtt EK505, we first showed that human SAMD9L was also restrictive against this SIVcpzPtt strain. Yet, chimp and bonobo SAMD9Ls did not present an increased anti-SIVcpz activity compared to human SAMD9L, suggesting possible lentiviral-strain specificity. Therefore, chimp and bonobo SAMD9Ls seem to have an increased anti-HIV-1 activity, as compared to human SAMD9L. It is further possible that SIVcpz, naturally circulating in chimp populations, adapted to the *Pan* SAMD9L increased antiviral function.

### Bonobo-specific polymorphisms confer an increased anti-HIV-1 activity to human SAMD9L without compromising cellular protein synthesis

We wondered whether minimal changes in human SAMD9L informed from some of these natural *Pan* variants could impact human SAMD9L functions. We specifically tested the bonobo SNPs, which could constitute species-specific adaptations with functional implications. We therefore cloned L90S and/or K1446R variants in the context of the human SAMD9L plasmid and investigated their effects on two key functions of human SAMD9L: anti-lentiviral function as well as cellular protein synthesis shutdown (Fig. 6).

**Figure 6.**
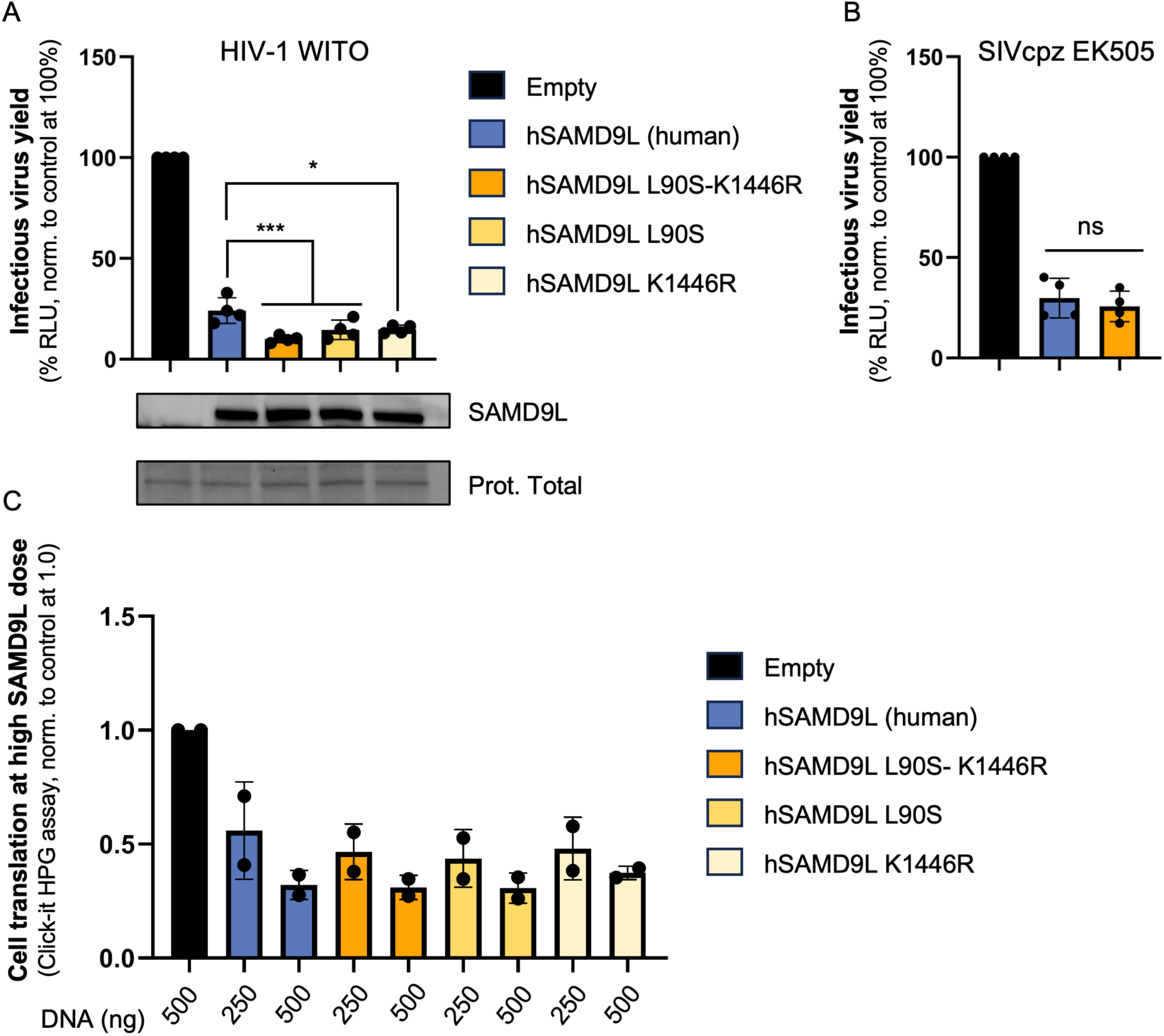
Bonobo-specific polymorphisms enhance human SAMD9L anti-HIV-1 activity, without affecting its translation shutdown effect. **A-B,** Relative infectious virus yields of HIV-1 pWITO (A) and SIVcpz EK505 (B) in indicated SAMD9L conditions (hSAMD9L, human SAMD9L), normalized to the empty control. Statistics were performed using the ratio paired t-test versus the human SAMD9L condition (***, p value <0.0005; *, p value < 0.05). Below, Western-blot analysis in the producer cells showing similar protein expression of SAMD9Ls. Loading control is from total protein (Prestained gels). In B, empty and hSAMD9L wt conditions are identical to Fig. 5E. **C,** Bonobo-specific SNPs do not modify human SAMD9L restriction of cellular translation. Quantification of protein synthesis assay was performed in two independent biological replicates in the context of ectopic expression of SAMD9Ls. Two doses of input DNA plasmids per condition were tested. HPG MFI ratio was calculated within each experimental condition using the MFI of the mCherry^+^ cells (expressing mCherry-SAMD9L) normalized to the MFI of the mCherry^-^ cells (not expressing SAMD9L). MFI, median fluorescence intensity.

First, we investigated the effect of these bonobo-specific variants on human SAMD9L function in the context of lentiviral replication, as in Fig. 5. Interestingly, we found that ectopic SAMD9L-L90S/K1446R and the single variants were expressed at a similar level to WT SAMD9L, but had a significant two-fold increase in anti-HIV-1 pWITO activity (Fig. 6A). The double mutant SAMD9L-L90S/K1446R appeared the most restrictive, while the single mutants had intermediate effects, suggesting additive functions (Fig. 6A). The increased effects of the human SAMD9L mutant with L90S and/or K1446R seemed independent of HIV-1 protein translation shutdown (Fig. S6A).

Second, we tested the activity of SAMD9L-L90S/K1446R on SIVcpzPtt EK505 replication and found that, similarly to wt *Pan* SAMD9Ls, the specific-SNPs in the context of the human SAMD9L did not increase its anti-SIVcpz function (Fig. 6B).

Lastly, we assessed the extent of whole cellular translation in human 293T cells transfected with high doses of either wild-type (WT) human mCherry-SAMD9L or mCherry-SAMD9L-L90S/K1446R or the single mutants. We used Click-iT™ L-Homopropargylglycine (HPG) Synthesis Assays, which measures the incorporation of HPG an analog of Methionine into newly synthesized proteins by flow cytometry. As previously shown ^5,8,61,62^, we found a dose-dependent shutdown of cellular translation in cells ectopically expressing human SAMD9L (mCherry^+^) normalized to the control mCherry^-^ cells (Fig. 6C). Importantly, we found that hSAMD9L-L90S/K1446R and the single variants displayed no differences in the cellular translation repression compared to the human WT hSAMD9L (Fig. 6C). This suggests that the non-synonymous bonobo variants in SAM and TPR domains do not change the activity of human SAMD9L on cellular protein synthesis.

Overall, bonobo-specific polymorphisms specifically enhance human SAMD9L antiviral function against HIV-1, without affecting the translation shutdown function.

## Discussion

This study unveils key aspects of the functional evolution of SAMD9s at different time scales, highlighting multidomain and functional convergence between metazoan SAMD9s and prokaryotic Avs9s, as well as recurrent genetic and genomic adaptations in mammals from ancient to very recent times. On one hand, we identified SAMD9/9L structural analogs in bacterial defense systems that induce cell death with similar AlbA_2 effector determinants, suggesting a remarkable shared immune factor across billions of years. On the other hand, our analyses of mammalian SAMD9/9L revealed dynamic and episodic adaptations, notably in primates, probably in response to epidemics, including from lentiviruses. By an “immuno-evo” framework, we bring key insights to understand the duality between the maintenance of a key antiviral shared immunity and its constant adaptations to pathogens.

### SAMD9/9L as a shared ancestral antiviral immune defense

The *SAMD9* gene family may be part of a shared ancestral immunity, in which eukaryotic antiviral mechanisms were acquired from bacteria by horizontal gene transfer, by vertical inheritance originating from LUCA (Last Universal Common Ancestor), or from convergent evolution ^16,18,66^. The list of ancestral immune defense systems is rapidly expanding with currently about a dozen identified antiviral systems, including cGAS, Viperin and TLRs^18^. Biochemistry, mechanistic and biochemical studies in both eukaryotic and prokaryotic systems will enable advancement into understanding these major immune systems. Here, we revealed striking structural similarity between human antiviral SAMD9/9L and prokaryotic Avs proteins ^37^, specifically with a new Avs protein family (Avs9). Notably, they both share the key nuclease domain, AlbA_2, which, in human SAMD9/9L, is responsible for tRNA^Phe^ cleavage and viral and cellular translational inhibition^5,9^. In Avs9 from *Pseudomonas f.*, we uncovered cell killing activity, which also depended on AlbA_2 and its predicted nuclease site. Future biochemical and functional investigations will determine if Avs9 has *bone fide* nuclease activity – and which substrate, as well as identify if Avs9 has specific anti-phage functions. Furthermore, similar to the Avs system, in which infected bacteria employ an altruistic self-killing mechanism for the benefit of the colony ^37^, tight regulation of SAMD9/9L is likely essential and is certainly possible thanks to the intermediate and C-terminal domains ^8^.

Uncovering phylogenetic relationships of prokaryotic Avs9 and metazoan SAMD9 families strongly suggest that both result from evolutionary convergence. Precisely, our study exemplifies how advantageous multi-domain combinations can arise independently through convergent evolution. Domain shuffling is a major driver of protein evolution and can lead, under analogous evolutive pressures, to similar multidomain proteins, resulting in surprising evolutionary and functional relationships across extremely long timescales. The convergence of Avs9 and SAMD9 is a remarkable example of complex multi-domain protein evolution between bacteria and humans.

### Extensive, and likely adaptative, *SAMD9/9L* copy number variations

Despite its simultaneous presence in evolutionarily distant organisms, we globally found a patchy distribution of *SAMD9* homologs and structural analogs across domains of life. For example, plants, fungi, algae and protozoa do not seem to harbor complete SAMD9 homologs. Furthermore, we found a very rapid evolution at the genomic and genetic levels during mammalian evolution in SAMD9/9L. Therefore, this ancient conservation is concomitant with rapid evolution, almost certainly as the result of virus-host arms-races. Gene loss and duplication may, for example, provide an advantage, similar to observations in other innate immune genes, like the APOBEC3 family ^31,67–69^ and many other factors^1^. It is noteworthy that most genomic variations were gene losses rather than extensive duplications, at least in most analyzed mammalian species.

At the mechanistic level, while the majority of *SAMD9/9L* losses occurred by genomic loss of a chromosomal region, *Colobinae* primates seem to have lost *SAMD9* by early stop codons and pseudogenization. However, it cannot be excluded that, in some species, this may lead to the expression of a truncated SAMD9 retaining the AlbA_2 effector function but lacking the crucial regulatory intermediate and C-terminal domains, similar to some SAMD9L autoinflammatory gain-of-function variants for example ^8,61^. Further study on its selective pressure, and on mRNA transcript and protein expression in natural tissues from *Colobinae* species under immune stimulation would resolve this question.

### The specific case of *SAMD9* unfixed loss in bonobos and adaptive chimp and bonobo SAMD9Ls: modern implications, functions, and potential past lentiviral drivers

Bonobos harbor a recent unfixed loss of *SAMD9*, which occurred through a large chromosomal deletion. Despite bonobos, chimps and humans being closely related, the variability observed at this locus is intriguing. Two bonobo-specific missense polymorphisms in SAMD9L that confer an increased antiviral activity against HIV-1 are located in the SAM (Sterile alpha motif) and TPR (Tetratricopeptide repeat) domains, which could be involved in protein-protein or protein-RNA interactions ^37,70–72^. One possibility is that the variants may modulate SAMD9L sensing, especially for the TPR domains, which have been reported to act as (viral) sensors in Avs and IFIT proteins (IFN-induced protein with tetratricopeptide repeats) ^37,73^. Further, although most human deleterious gain-of-function mutations in SAMD9/9L associated diseases are in the P-loop NTPase domain, some are described in the TPR or SAM domains ^74^. However, we did not observe a gain-of-function phenotype on global cellular translation for the bonobo-specific variants. This therefore suggests that the variants may not destabilize the inactive closed form of the protein, nor change SAMD9L basal activities. Instead, it might modify its specificity and sensitivity in viral sensing, potentially adapting its interface with viruses. Otherwise, it may impact other potential functions of SAMD9L, for example in endosomal trafficking, increasing specific anti-HIV functions ^5,10,11^. It would also be interesting to determine if, and how, chimp and bonobo SAMD9Ls restrict other infections from poxviruses or other RNA viruses ^3,12^, and whether those may have driven some of the adaptations.

Our data show that chimp and bonobo SAMD9Ls have an increased anti-HIV-1 phenotype comparing to human. The additional loss of the pro-HIV-1 SAMD9^5^ may have been particularly advantageous (overall increased fitness) during past lentiviral infections in bonobo ancestors (i.e. increased antiviral SAMD9L and loss of prolentiviral SAMD9). The presence of 3/13 unrelated bonobos with one *SAMD9/SAMD9L* allele and one *SAMD9L*-only allele suggests that *SAMD9* loss is unfixed, and likely recent. If selection favors individuals without *SAMD9*, the gene could eventually be completely lost in the bonobo population. The modern genomic makeup of the *SAMD9/9L* locus in chimps, and even more so in bonobos, therefore suggests adaptation to lentiviral-like epidemics that occurred in *Pan*, as well as since the bonobo-chimp divergence. Performing this evo-functional study with a larger bonobo sample size would be necessary to determine the exact selective pressures shaping antiviral defense mechanisms in this species. Long-read sequencing of bonobos would also enable the determination of the haplotype structures on which the *SAMD9* gene was lost and the *SAMD9L* substitutions occurred, potentially providing insight into the epistatic interactions between these genetic changes.

Unlike some chimps and humans that are infected by SIVcpz and HIVs, respectively, and suffer from AIDS symptoms^75–77^, modern bonobos are not known to be naturally infected by any lentiviruses ^25,26^. Overall, SAMD9/9L adaptation may nowadays participate, with other factors ^78^, to bonobo population resistance against lentiviral/SIV infections.

Finaly, it is noteworthy that in this study SIVcpzPtt EK505 did not show an increased sensitivity to chimp and bonobo SAMD9Ls, or to *Pan* SNPs in the context of human SAMD9L. This may be the result of virus-host co-evolution^1,30^, where SIVcpz has adapted to the natural genetic makeup of its host, particularly of chimp antiviral innate immunity.

Altogether, our findings highlight the strength of evo-immuno approaches in unraveling links between the evolutionary history of innate immunity and contemporary challenges in human health. The identification of SAMD9/9L homologs and structural-functional analogs across diverse taxa, as prokaryotes and primates, shows a shared ancestral immunity. Common challenges, such as fighting viral infections, drive both conservation or convergence of key immune systems as well as their rapid evolution through arms-races. In this regard, using diverse models (human, diverse eukaryotic cells, and bacteria) and natural variants in closely related species for functional studies can bring valuable insights with broader medical applications, such as the incorporation of potentiator mutations in antiviral factors (protein engineering) or the use of bacterial antiviral proteins that could act against human viruses.

## Methods

### Comparative genomics, phylogenetics, and positive selection analyses in mammals

To obtain the coding sequences of the SAMD9 and SAMD9L homologs in bats, rodents, primates, ungulates and carnivores, we used the Detection of Genetic INNovations (DGINN) pipeline ^53^ with, respectively, *Myotis myotis, Rattus norvegicus, Homo sapiens, Hippopotamus amphibius*, and *Phoca vitulina* Refseq SAMD9 and SAMD9L, as queries. Briefly, the coding sequences from each group were automatically retrieved with NCBI blastn ^79,80^, cleaned, and aligned with MAFFT ^81^. Homologous sequences from marsupials, aves, and amphibians were retrieved using NCBI Blastn. Of note, these analyses are based on publicly available genome annotations (not necessarily genes annotated as “SAMD9/9L-like” but regions annotated as coding regions), so it is not excluded that some unannotated genes were not analyzed. The species and accession numbers are presented in Supplementary Table 1. Nucleotide alignments of SAMD9 or SAMD9L from each mammalian group were manually curated before being used as input in DGINN for automatic codon alignment using the Probabilistic Alignment Kit (PRANK) ^82^ and phylogenetic tree building using PhyML ^83^ (with default settings in DGINN).

Furthermore, all codon alignments, as well as outgroup sequences, were realigned in a three-step fashion to obtain a high-quality mammalian wide codon alignment (i) using Muscle ^84^, (ii) manually curating the sequences, (iii) codon aligned with PRANK. A phylogenetic tree was inferred from this alignment using IQ-TREE web server (GTR+F+I+G4 identified as the best substitution model by ModelFinder implemented in IQ-TREE) ^85^. We also performed analyses with PhyML (best model estimated from Smart model selection SMS: GTR+R). These analyses allowed us to attribute the phylogenetically-aware “SAMD9” or “SAMD9L” nomenclature.

To test for branch-specific, episodic diversifying selection, the codon-alignments of primate SAMD9 and SAMD9L with *Cricetulus griseus* (criGri) and/or *Tupaia chinensis* (tupChi) as outgroups were used as inputs for the adaptive branch-site random effects likelihood (aBSREL) program on the DataMonkey webserver ^57^. Each branch that we tested for evidence of positive selection was defined as “tested branch” (i.e. “foreground”) and the remaining as “background branches”. Two models were fit to each tested branch: one that allows for episodic diversifying selection (with an ω > 1), and one that does not. A likelihood ratio test is then used to compare these models and assess whether the tested branch shows evidence of positive selection. For branches where selection is detected, aBSREL also estimates the proportion of codon sites that are subject to positive selection.

For cases in which we suspected gene losses, we confirmed the absence of coding genes by several methods. We analyzed the SAMD9/9L genomic locus (between *HEPACAM2* et *CDK6* syntenic genes) on NCBI genome data viewer of specific species. We verified that there were no missing data or low sequence quality in this genomic region. Pseudogenes, here identified by multiple and very early stop codons, were only analyzed systematically in primates. For the two monotreme species, in which no SAMD9/9L homologs could be retrieved by genome-wide blast, the HEPACAM2-CDK6 genomic locus was retrieved. We found no missing data (no “N”) and no homology using blast or alignments (with relaxed parameters) of non-annotated regions with human SAMD9/9L.

### Genome alignment and single nucleotide polymorphisms analyses in hominids

The genomic sequences of thirteen bonobos, 59 chimpanzees and 4099 humans were retrieved from public online databases: 1000 Genomes Project (1kGP), Human Genome Diversity Project (HGDP), NCBI bioprojects PRJNA189439, SRP018689, PRJEB15083 ^60,60,86^ (The 1000 Genomes Project Consortium 2015). DNA sequences from *Pan* individuals were aligned to Clint_PTRv2/panTro6 reference genome using BWA-MEM ^87^. Then, variant calling was done using FreeBayes ^88^ to obtain a vcf file. The SAMD9 and SAMD9L locus regions (chr7:88599930-89350079 in panTro6) were extracted and parsed using an *ad-hoc* R script using VariantAnnotation, GenomicFeatures, AnnotationHub, org.Pt.eg.db, ggplot2, R packages. This script was used to identify non-synonymous variants among SNPs and to visualize them. The equivalent genomic region in human was retrieved by using the LiftOver tool of the UCSC genome browser (chr7:92540798-93289621 in the human reference GRCh38/hg38). These coordinates were then used to subset the HGDP+1000 Genomes Project vcf for this region.

### Structure similarity search, structurally-aware alignment, and phylogenetic analyses across kingdoms of life

Protein structures were obtained from AlphaFold DB ^89,90^ and RCSB PDB ^91^. Foldseek ^33^ was employed for detection of structural similarity. We used a sequential strategy. First, we queried Foldseek v.427df8a with SAMD9 (PDB ID: Q5K651) and SAMD9L (Q8IVG5) against the AlphaFold database clustered at 50% sequence identity (AFDB50), using a 30% TM-score threshold and a maximum E-value of 0.0001 (default settings). Then, we constructed a query database consisting of SAMD9 (Q5K651), SAMD9L (Q8IVG5), and bacterial top hits — AVAST type V (A0A1F9N8W4) and Avs9 (A0A100VJR7, A0A2S4Y961 and A0A7T4VS34) — and used it for searches under the same settings. Search hits were subsequently filtered for a minimum 80% query coverage to cover at least 1000 aa of the query structures, resulting in 238 analog structures. FoldMason v.333d54c ^92^ was used to generate a multiple structure alignment (MSTA) of the identified structural analogs and MUSCLE v5 was used to generate a multiple amino acid sequence alignment (MSA). The domain coordinates used for presence/absence analysis of each domain or for the extraction of a given domain MSTA are presented in Fig. S1C and are based on the predicted SAMD9 3D structure. Additionally, using FoldMason, a MSTA was generated on the human SLFNs (SLFNL1, SLFN5, SLFN11-14, respectively corresponding to PDB IDs Q499Z3, Q08AF3, Q7Z7L1, Q8IYM2, Q68D06 and P0C7P3) with the 23 hits over 238 containing an AlbA_2 domain. Phylogenetic trees were constructed from both the MSA and MSTA using IQ-TREE 2.3.0 ^93^ with the LG+F+G4 substitution model, 1000 bootstrap replicates, and visualized using the ggtree R library ^94^ or iTol ^95^.

### Phylogenetic analysis of prokaryotic and eukaryotic AlbA_2 domains

The HMM profile of the Pfam AlbA_2 (PF04326) domain was retrieved from the Pfam database ^96^. This profile was searched against a custom protein database combining : i) 41,150 complete bacterial genomes downloaded from Refseq in August 2024, filtered for redundancy using the clusthash function of MMseqs2 (v 13.45111) using default parameters ^97^; ii) 455 complete archaeal genomes downloaded from Refseq in August 2024; iii) 993 representative eukaryotic genomes from the EukProt database ^98^, filtered for redundancy using the clusthash function of MMseqs2 (13.45111) using default parameters. The AlbA_2 HMM profile was searched into this combined protein database using hmmsearch (v3.3.2) with default parameters ^99^. Hits with at least 90 covered profile residues were selected and the amino acid sequences of the aligned regions complemented with ten residues on each side were extracted. Sequences were clustered using the easy-cluster function of MMseqs2 (v13.45111) with parameter --min-seq-id 0.8 ^97^. Cluster representatives were aligned with Clustal-Omega ^100^ with default parameters and the alignment was trimmed using ClipKit ^101^. The trimmed alignment was used to compute a tree using IQ-TREE with parameters -m L -bb 10000 -nm 10000, which was then visualized using iTOL.

For each genome, defense systems were detected using DefenseFinder (v 1.3)^34^. For each clade, we calculated a defense score as the fraction of bacterial genes found within 10 genes upstream or downstream of a defense protein as detected by DefenseFinder in their genome of origin.

### Bacterial strains and plasmids

The codon-optimized open reading frame encoding Avs9 from *P. fluorescens* (Uniprot ID A0A7T4VS34) was ordered as a gene fragment from Twist Bioscience, cloned into the pBbA6c vector ^102^ by T5 exonuclease-dependent assembly (TEDA) cloning ^103^ and transformed into *E. coli* DH5α λpir. Constructions were sequenced-verified by Sanger sequencing (Microsynth). The complete Avs9 gene fragment sequence synthesized and used in this study is available (see Data Availability). We further made the Avs9-D45N mutant by site directed mutagenesis using PCR amplification with Q5 polymerase (New England Biolabs) and KLD cloning (New England Biolabs).

### Bacterial drop assays

*E. coli* DH5α λpir cells carrying pBbA6c, pBbA6c-Avs9 or pBbA6c-Avs^D45N^ were grown for 6 h at 37°C and 180 rpm in Luria-Bertani (LB) medium supplemented with chloramphenicol (Cm) 20 μg/ml and glucose 1 g/mL. After 10-fold serial dilutions, 5 µL of each dilution were spotted on LB agar plates supplemented with 20 µg/mL Cm and either 1% glucose or 100 to 500 µM Isopropyl β-D-1-thiogalactopyranoside (IPTG). Plates were incubated at 37°C and 25°C for 24 h and 48 h respectively. Strong toxicity of pBbA6c-Avs9 was observed upon incubation at 25°C.

### Plasmids for expression in human cells

HIV-1 T/F pWITO (Human Immunodeficiency Virus 1 pWITO.c/2474, ARP-11739) encoding a full-length infectious molecular clone (IMC) was contributed by Dr. John Kappes and Dr. Christina Ochsenbauer through the NIH AIDS repository program. IMC for SIVcpzEK505 was a gift from Beatrice Hahn^27,65^. The pMT06-Flag-mCherry-SAMD9L plasmid was constructed by cloning the synthetized human *SAMD9L* gene into a pMT06-Flag-mCherry backbone, from the original RRL.sin.cPPT.CMV/Flag-E2-crimson.IRES-puro.WPRE (MT06, a gift from Caroline Goujon: Addgene plasmid # 139448; http://n2t.net/addgene:139448; RRID:Addgene_139448)^104^. The pMT06-Flag-mCherry-chimpSAMD9L plasmid (chimpSAMD9L) was synthesized and cloned by Azenta Genewiz. The bonobo-SAMD9L plasmids were generated through site-directed mutagenesis of the pMT06-FLAG-mCherry-chimpSAMD9L plasmid. Human SAMD9L-L90S/K1446R, SAMD9L-L90S and SAMD9L-K1446R (double and single mutant) plasmids were generated from pMT06-Flag-mCherry-SAMD9L plasmid, using the QuikChange Lightning Site-Directed Mutagenesis Kit (Agilent) following the manufacturer’s instructions. Sequences were confirmed through full-length plasmid and/or Sanger sequencing (Microsynth).

### Cell lines and culture

Human embryonic kidney 293T (ATCC, cat. CRL-3216) and TZM-bl (NIH AIDS Research and Reference Reagent Program, Cat. 8129) cell lines were grown in Dulbecco Modified Eagle Medium (DMEM) containing 10% fetal calf serum (FCS, Sigma cat. F7524) and 100 U/ml of penicillin/streptomycin. TZM-bl cells express the cell surface proteins CD4, CCR5 and CXCR4, and encode for Luciferase and β-galactosidase under the control of the LTR promoter. They are commonly used for lentiviral titration of culture supernatants (Tzm-bl assays).

### Production and quantification of replication-competent lentivirus

293T cells were initially seeded in 6-well plates at a density of 0.2M cells/ml (400,000 cells total per well). After 24 hours, the cells were co-transfected using TransIT-LT1 (Mirus) with a plasmid encoding a fully replication-competent lentivirus (IMCs), alongside either a plasmid encoding SAMD9L or an empty control. The quantity of DNA used was 250 ng for host plasmids and 1200 ng for virus plasmids. Subsequently, 48 hours post-transfection, cells were harvested for Western blot. The supernatants were collected and stored at -80°C for further titration of infectious virus yield via TZM-bl cells. For titration, TZM-bl cells were plated in 96-well plates and exposed to serial dilutions of viral supernatant. Following 48 hours of infection, cell lysis was performed using BrightGlow Lysis Reagent (Promega E2620), and relative light units (RLU) were measured using the Tecan Spark® Luminometer. Infectious virus yields under various conditions were consistently expressed as fold-change compared to paired viral infection conditions in the absence of SAMD9L.

### Western blot analysis

Cells were harvested and lysed using ice-cold RIPA buffer (composed of 50 mM Tris pH8, 150 mM NaCl, 2 mM EDTA, and 0.5% NP40) supplemented with protease inhibitors (Roche), followed by sonication. Proteins from cell lysates or supernatants were separated by electrophoresis and transferred onto a PVDF membrane via overnight wet transfer at 4°C. Stain-Free gel (BioRad) was used for loading and protein transfer controls. Following blocking in TBS-T 1X solution (Tris Buffer Saline, consisting of Tris HCl 50 mM pH8, NaCl 30 mM, and 0.05% Tween 20) with 5% powdered milk, the membranes underwent incubation with primary antibodies for a duration ranging from 1 hour to overnight, followed by subsequent 1-hour incubation with secondary antibodies. Detection was carried out using SuperSignal West Pico Chemiluminescent Substrate (ThermoFisher Scientific) and imaged using the Chemidoc Imaging System (BioRad). Antibodies utilized included anti-SAMD9L (Proteintech, 25173-1-AP), anti-Gag (NIH HIV Reagent Program, 183-H12-5C), anti-HIV-1-gp120 (Aalto, D7324; NIH HIV Reagent Program, 16H3), as well as secondary IgG-Peroxidase conjugated anti-mouse (Sigma, cat. A9044) and anti-rabbit (Sigma, cat. AP188P). “Total protein” was used as a loading control with BioRad Stain-Free gel.

### Protein synthesis assay

293T cells were seeded at 0.2M cells/ml in 12-well plates (200,000 cells total per well). Twenty-four hours after seeding, cells were transfected with 250ng or 500ng of host DNA plasmid, using TransIT-LT1 (Mirus) following the manufacturer instructions. Forty-eight hours post-transfection, cells were incubated in L-homopropargylglycine (HPG) for 30 min at 37°C. Medium was discarded, and cells were harvested and fixed with PFA 4%. Cells were then washed with PBS BSA 3% and permeabilized in PBS 0,5% Triton X-100 for 15 min. Click-iT® Plus Alexa Fluor® Picolyl Azide assay was then performed and cells were analyzed on MACSQuant® VYB Cytometer (Miltenyi Biotec, SFR BioSciences).

### Other Softwares and Statistical Analyses

DefenseFinder ^34^was used to analyze sequences from Foldseek analyses ^33^. Sequencing analyses and representations were conducted using Geneious (Biomatters), ESPript 3.0 https://espript.ibcp.fr ^105^ and UGENE v52.0 ^106^. R scripts were used to conduct analyses of genomic data. Graphic representations and statistical analyses were carried out using GraphPad Prism 9 and R scripts. In the figures, data are presented as mean ± SD, and each point correspond to an independent biological replicate. Statistics were performed using the ratio paired t-test (*, p value < 0.05; **, p value < 0.005).

### Availability of codes, data and reagents

All data and reagents are available in this manuscript (including in Dataset S1-S3) or accessible upon request to the corresponding author. The scripts for the analyses of the polymorphic sites are available at http://gitbio.ens-lyon.fr/ciri/lp2l/vcf_to_nice_figures.git. The alignments and phylogenetic trees are all openly available through FigShare deposits (Phylogenetics associated: with Fig. 1 and S1 https://doi.org/10.6084/m9.figshare.29082653 ; with Fig. 2 and S2 https://doi.org/10.6084/m9.figshare.29082710 ; with Fig. 3 and S3 https://doi.org/10.6084/m9.figshare.29042762 ; with Fig. 4 and S4 https://doi.org/10.6084/m9.figshare.29087987 ; with Fig. S5C https://doi.org/10.6084/m9.figshare.29088050). Avs9 sequence used for experimental assays in Fig. 3D and S3D is available at https://doi.org/10.6084/m9.figshare.29082719.

## Supporting information

Datase S2

## Acknowledgements

We sincerely thank Francesca Fiorini (MMSB), Florence Roucher-Boulez (HCL), Aude Bernheim and Hugo Vaysset (Pasteur Institute) for feedback on the project, LP2L past and present lab members for support and discussion, Laurent Guéguen (LBBE) for providing guidance in the positive selection analyses in DGINN. We thank Amandine Le Corf (CIRI), Caroline Goujon (IRIM), Laura Guiguettaz and Emiliano Ricci (LBMC), Beatrice Hahn (Univ. of Pennsylvania, Perelman School of Medicine), Michael Emerman (Fred Hutch) and the NIH AIDS reagent program for sharing reagents. We acknowledge the contribution of SFR Biosciences (Université Claude Bernard Lyon 1, CNRS UAR3444, Inserm US8, ENS de Lyon) ANIRA platform, especially Véronique Barateau for her assistance, as well as Didier Decimo of the BSL-3 ENS-Lyon platform. We also thank all the contributors of the publicly available bioinformatic programs and genomic sequences.

## Author contributions (CRediT Contributor Roles Taxonomy)

**Author contributions (CRediT: Contributor Roles Taxonomy)**
|  |  |
| --- | --- |
| Conceptualization | Ideas; formulation or evolution of overarching research goals and aims<br>AL, LE |
| Methodology | Development or design of methodology; creation of models<br>AL, ACh, RD, CLMG, MS, FR, PHS, LE |
| Software | Programming, software development; designing computer programs; implementation of the computer code and supporting algorithms; testing of existing code components<br>AL, ACh, CR, CLMG, JLR, MS, FR, PHS, LE |
| Validation | Verification, whether as a part of the activity or separate, of the overall replication/ reproducibility of results/experiments and other research outputs<br>AL, ACh, RD, CL, FR, LE |
| Formal analysis | Application of statistical, mathematical, computational, or other formal techniques to analyze or synthesize study data<br>AL, ACh, RD, CLMG, JLR, LP, MS, FR, PHS, LE |
| Investigation | Conducting a research and investigation process, specifically performing the experiments, or data/evidence collection<br>AL, ACh, RD, CR, JB, CLMG, JLR, CL, LP, FR, PHS, LE |
| Resources | Provision of study materials, reagents, materials, patients, laboratory samples, animals, instrumentation, computing resources, or other analysis tools<br>AL, ACh, RD, CR, CLMG, AC, MS, FR, PHS, LE |
| Data Curation | Management activities to annotate (produce metadata), scrub data and maintain research data (including software code, where it is necessary for interpreting the data itself) for initial use and later reuse<br>AL, ACh, MS, CLMG, FR, PHS, LE |
| Writing - Original Draft | Preparation, creation and/or presentation of the published work, specifically writing the initial draft (including substantive translation)<br>AL, ACh, LE |
| Writing - Review & Editing | Preparation, creation and/or presentation of the published work by those from the original research group, specifically critical review, commentary or revision – including pre- or postpublication stages<br>All authors : AL, ACh, RD, CR, JB, CLMG, JLR, CL, LP, AC, MS, FR, PHS, LE |
| Visualization | Preparation, creation and/or presentation of the published work, specifically visualization/ data presentation<br>AL, ACh, RD, CR, CLMG, JLR, FR, PHS, LE |
| Supervision | Oversight and leadership responsibility for the research activity planning and execution, including mentorship external to the core team<br>FR, PHS, LE |
| Project administration | Management and coordination responsibility for the research activity planning and execution<br>LE |
| Funding acquisition | Acquisition of the financial support for the project leading to this publication<br>AL, FR, PHS, LE |

## Supplementary Figures

**Figure S1.**
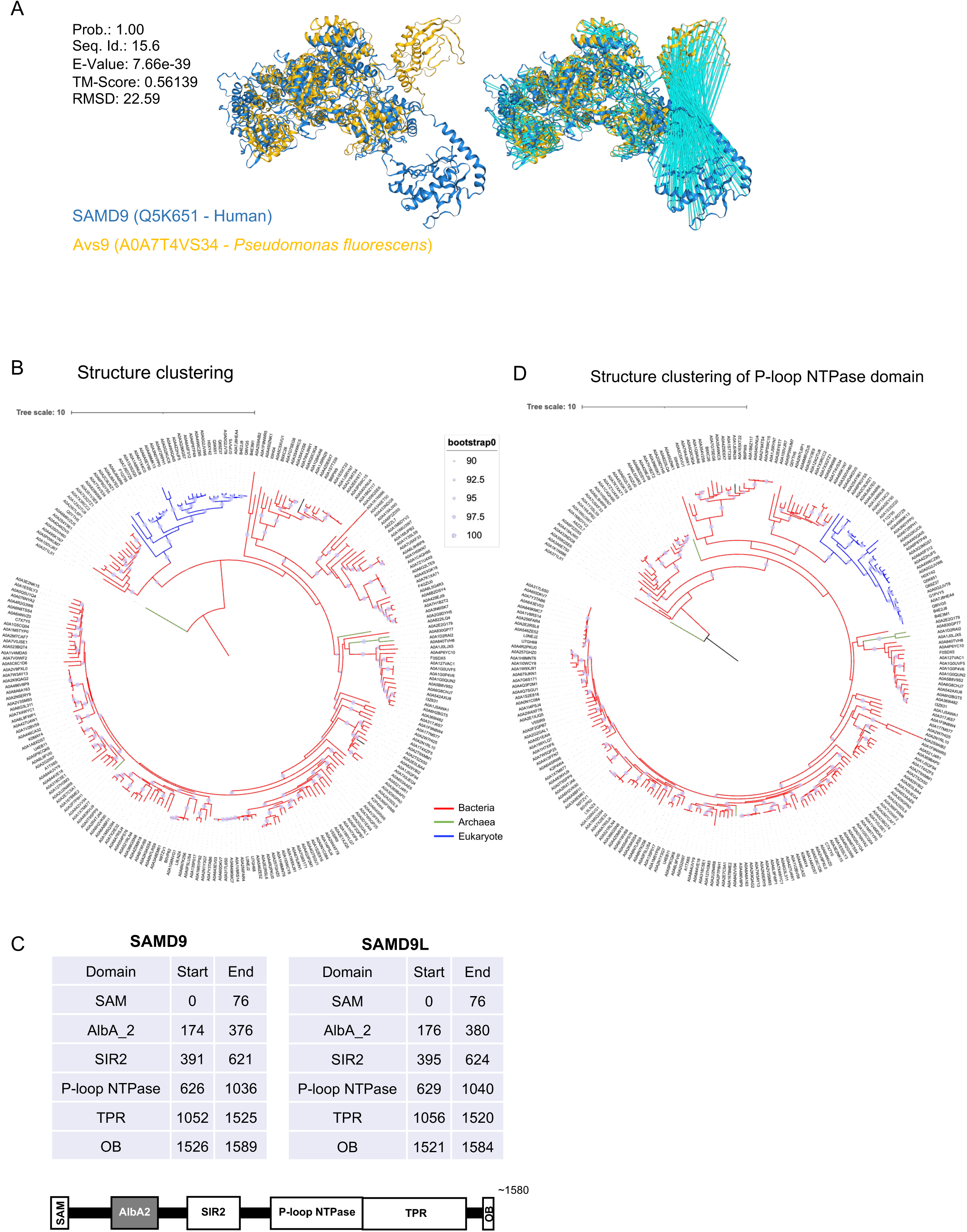
Associated to Figure 1. SAMD9/9L structural analogs are extensively present in prokaryotes. **A,** Left. Protein overlap (FoldSeek) between human SAMD9 and *Pseudomonas fluorescens* A0A7T4VS34 (Avs9). Right. Similar with blue bars linking aligned residues. **B,** Structure clustering tree shows widespread and diverse SAMD9/9L analogs in bacteria. **C,** P-loop NTPase structure clustering tree. **D,** SAMD9/9L domain bounds (coordinates of the protein sequence) used for the biocomputational analyses.

**Figure S2.**
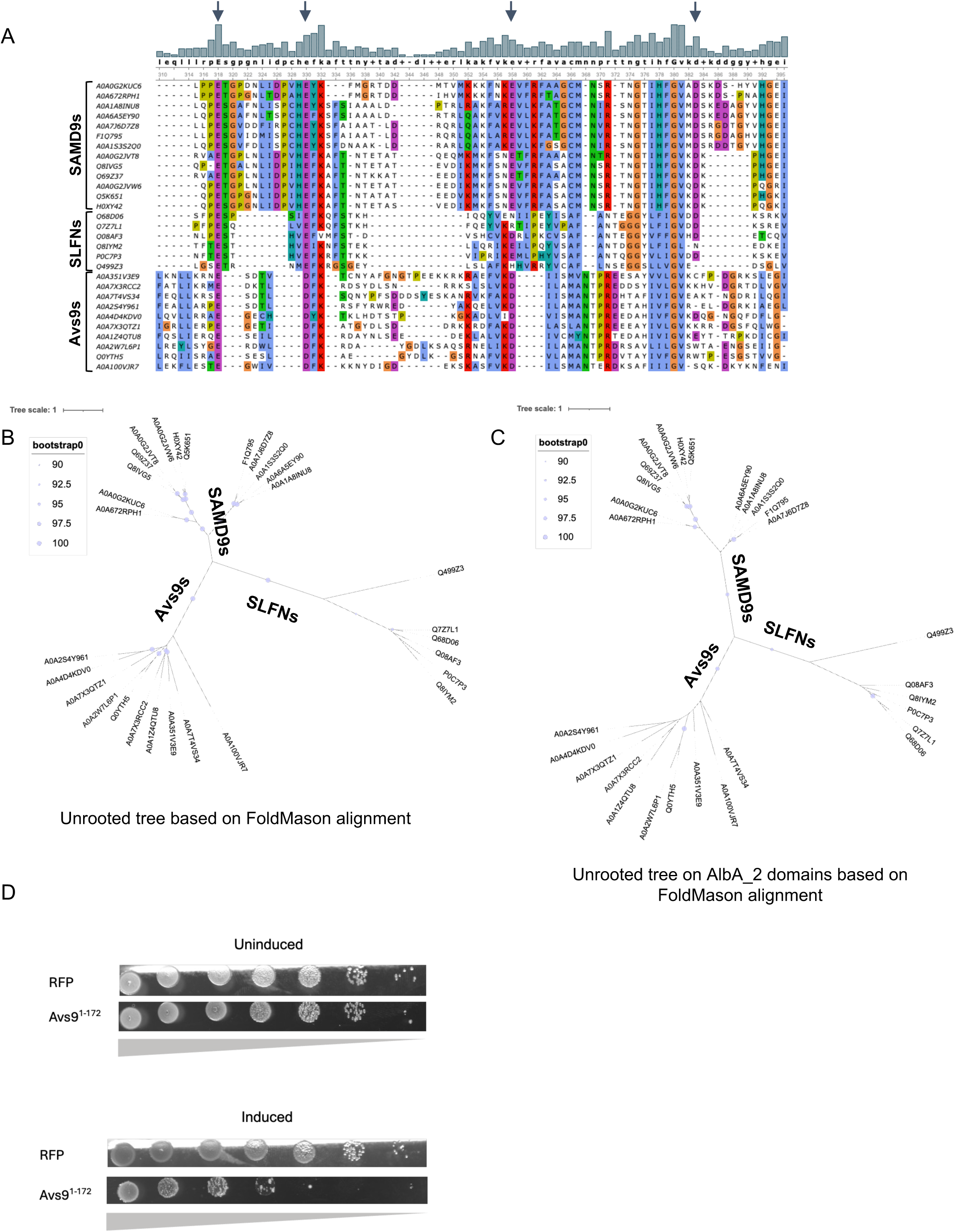
Associated to Figure 2. AlbA_2 effector domain is shared between human SLFNs, SAMD9s and bacterial Avs9, with a conserved catalytic site (A-C), and AlbA_2 effector domain of Avs9 is sufficient for the cell death induction in bacteria (D). **A,** Representation of the amino acid sequence alignment (from Muscle) showing the shared conserved residues (highlighted by the arrows) between the different protein families – SLFNs, SAMD9s and Avs9s. The alignment was performed and displayed on UGENE v52.0 with Clustal X colors. **B-C,** Structure clustering of full-length protein (B) or AlbA_2 domain extraction (C) trees shows three distinct clades for SLFNs, SAMD9s and Avs9s proteins respectively. **D,** Bacterial drop assay of bacteria transformed with either Avs9^1–172^ or RFP as a control, with induction (500µM IPTG) or without induction (1% glucose). Photo of serial dilutions of bacterial drops.

**Figure S3.**
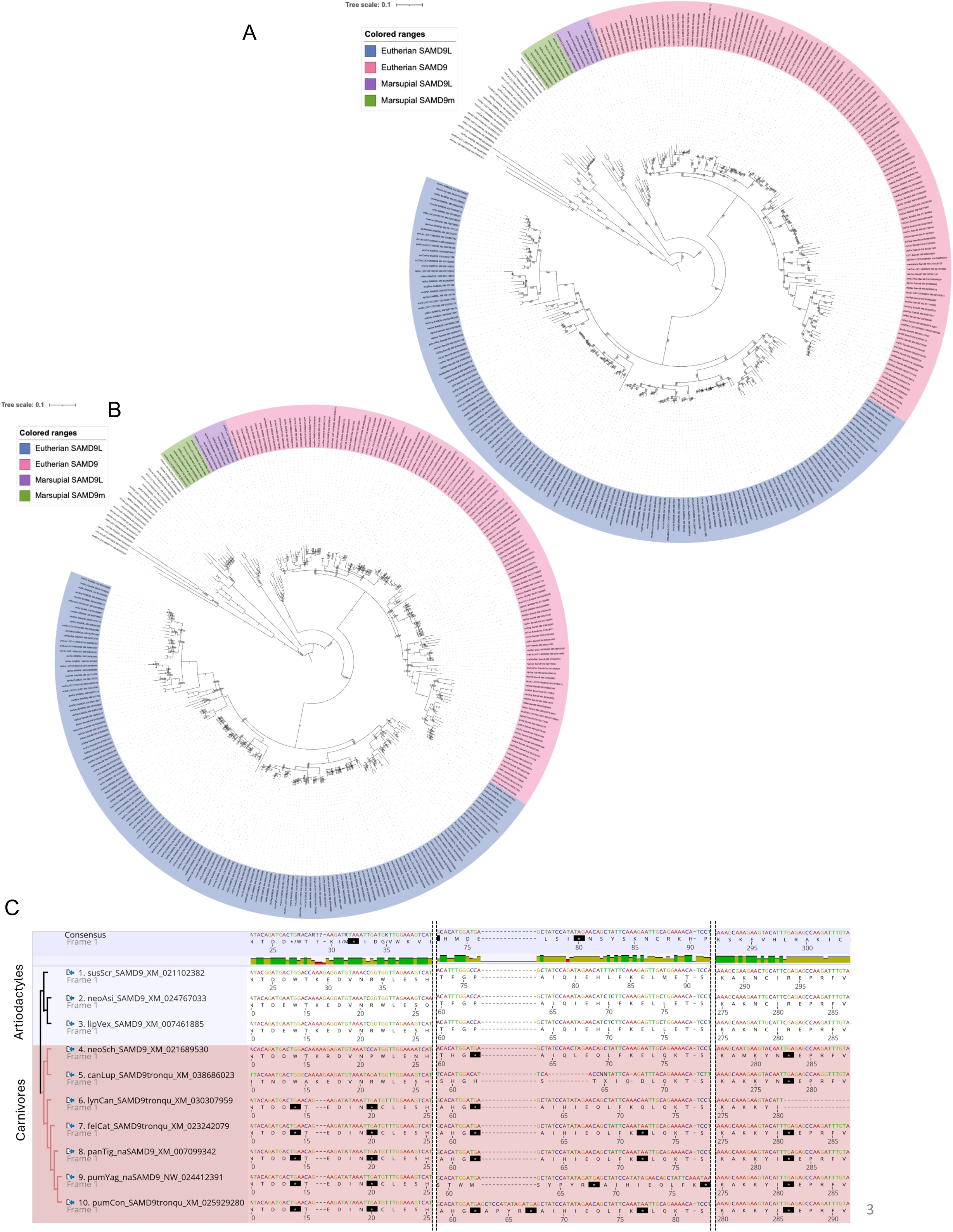
Associated to Figure 3. Phylogenetic analyses of SAMD9/9L in mammals associated with Figure 3. **A,** Maximum likelihood phylogenetic tree of all vertebrate SAMD9s from Dataset S1, determined using IQ-TREE web server (GTR+F+I+G4 identified as the best substitution model by ModelFinder implemented in IQ-TREE). Ultrafast bootstrap support values are shown next to branches. **B,** Maximum likelihood phylogenetic tree determined using PhyML (best model estimated from Smart model selection SMS: GTR+R). aLRT support values are shown next to branches. **C**, Parts of the codon alignment of representative artiodactyl and carnivore SAMD9s, highlighting early STOP (*) codons within carnivore SAMD9s. Amino acid (aa) numbering are shown below sequences. Snapshot from Geneious. Cladogram and sequence names shown on the left (species name abbreviated by three first letters of the genus and three first letters of the species (e.g. felCat, *Felis catus*).

**Figure S4.**
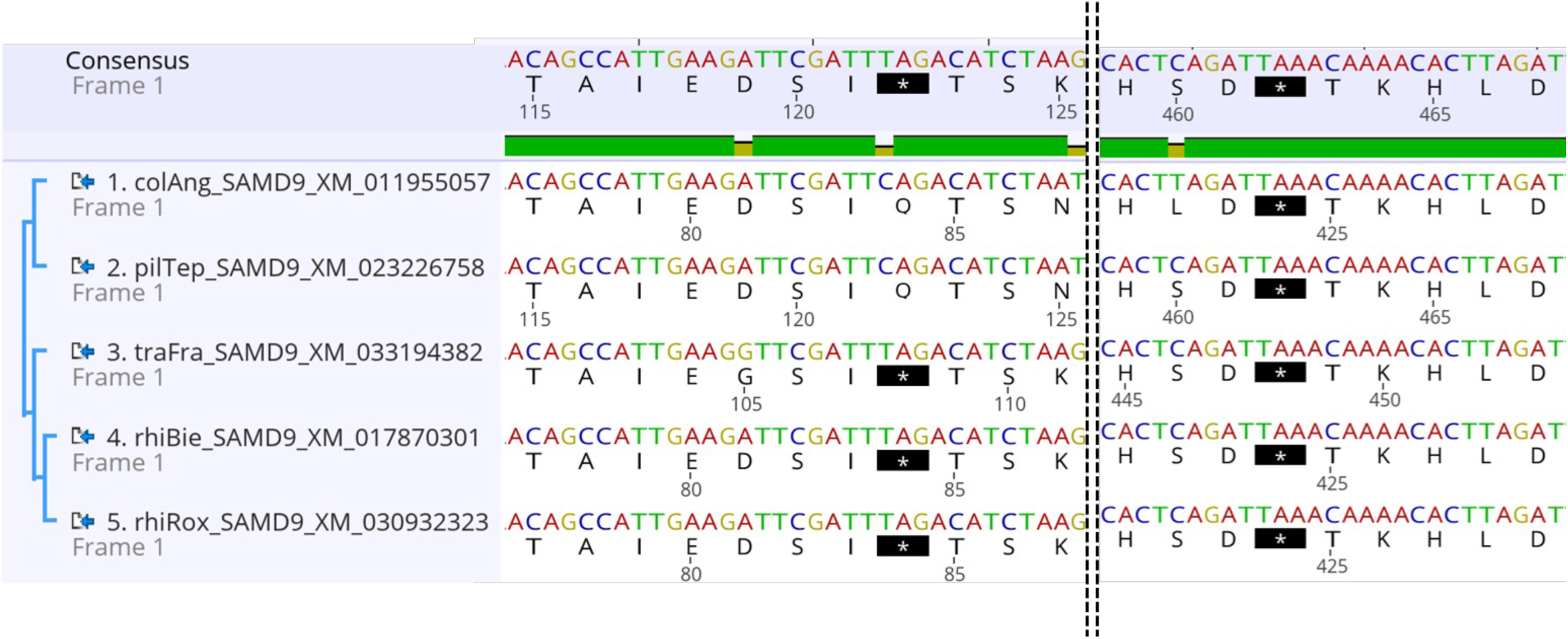
Associated to Figure 4. Evidence of early STOP codons in *Colobinae* SAMD9. Snapshot of parts of the codon alignment of *Colobinae* SAMD9 sequences with cladogram and sequence name shown on the left, highlighting early STOP (*) codons. Amino acid (aa) numbering is shown below each aa sequence. View from Geneious. Species name abbreviated by three first letters of the genus and three first letters of the species (e.g. colAng, *Colobus angolensis*).

**Figure S5.**
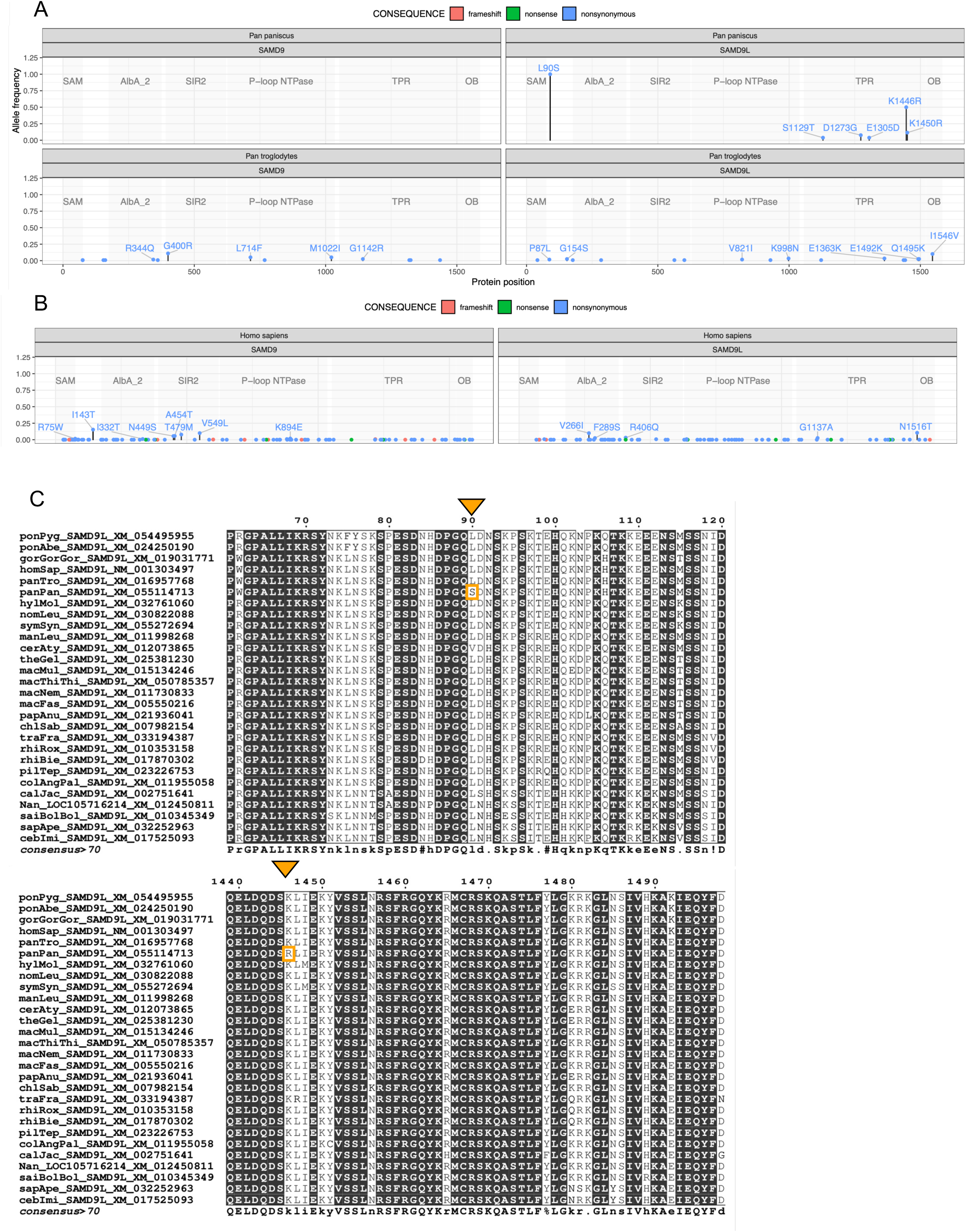
Associated to Figure 5. Alleles across SAMD9 and SAMD9L protein sequences in humans, bonobos and chimpanzees. **A-B,** Polymorphisms frequency in SAMD9 and SAMD9L among the *Pan* (A) and the human (B) populations, with PanTro6 and GRCh38/hg38 as references, respectively. The domains of SAMD9 and SAMD9L are delimited by transparent grey boxes. Labeled coordinates correspond to frequencies > 1%. **C,** Zoom on SAMD9L primate amino acid sequence alignment showing the bonobo-specific variants. Muscle aa alignment representation from ESPript 3.0, https://espript.ibcp.fr (Robert et Gouet 2014). Bonobo-specific variants are at position L90S and K1446R (highlighted in orange).

**Figure S6.**
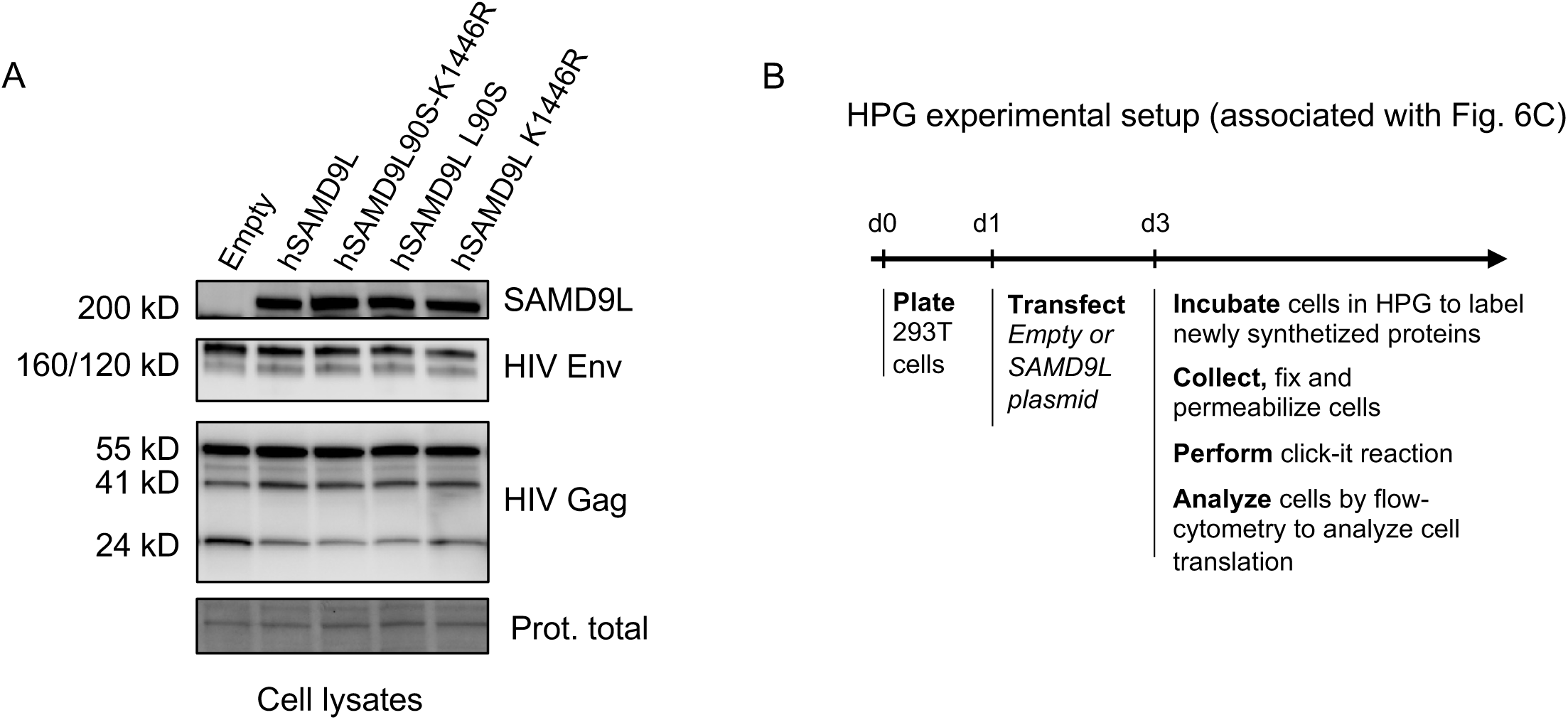
Associated to Figure 6. Effect of bonobo variants on human SAMD9L’s activities on cellular and viral protein translation. **A,** Western-blot analysis in the HIV-1 producer cells, following experimental setup of Fig. 5C and associated with experiment shown in Fig. 6A. Shown are the HIV-1 Gag (p55, p41, p24), HIV-1 Env (gp160, gp120), and SAMD9L protein expressions. Loading control is from total protein (BioRad Pre-stained gels). **B,** Experimental setup of the HPG assay to determine impact of SAMD9L bonobo SNPs on cell translation.

## Supplementary Dataset

**Dataset S1. References of sequences for vertebrate phylogenetic analyses.**

**Dataset S2. Gating strategy for flow cytometry experiments**

**Dataset S3. List of Foldseek hits (label, species, kingdom)**

## References

1. Tenthorey, J. L., Emerman, M. & Malik, H. S. Evolutionary Landscapes of Host-Virus Arms Races. Annu. Rev. Immunol. 40, 271–294 (2022).

2. Georjon, H. & Bernheim, A. The highly diverse antiphage defence systems of bacteria. Nat. Rev. Microbiol. 21, 686–700 (2023).

3. Liu, J. & McFadden, G. SAMD9 Is an Innate Antiviral Host Factor with Stress Response Properties That Can Be Antagonized by Poxviruses. J. Virol. 89, 1925–1931 (2014).

4. Meng, X. et al. A paralogous pair of mammalian host restriction factors form a critical host barrier against poxvirus infection. PLoS Pathog. 14, e1006884 (2018).

5. Legrand, A. et al. SAMD9L acts as an antiviral factor against HIV-1 and primate lentiviruses by restricting viral and cellular translation. PLOS Biol. 22, e3002696 (2024).

6. Meng, X. & Xiang, Y. RNA granules associated with SAMD9-mediated poxvirus restriction are similar to antiviral granules in composition but do not require TIA1 for poxvirus restriction. Virology 529, 16–22 (2019).

7. Sivan, G., Glushakow-Smith, S. G., Katsafanas, G. C., Americo, J. L. & Moss, B. Human Host Range Restriction of the Vaccinia Virus C7/K1 Double Deletion Mutant Is Mediated by an Atypical Mode of Translation Inhibition. J. Virol. 92, e01329–18, /jvi/92/23/e01329-18.atom (2018).

8. Russell, A. J. et al. SAMD9L autoinflammatory or ataxia pancytopenia disease mutations activate cell-autonomous translational repression. Proc. Natl. Acad. Sci. U. S. A. 118, e2110190118 (2021).

9. Zhang, F. et al. Human SAMD9 is a poxvirus-activatable anticodon nuclease inhibiting codon-specific protein synthesis. Sci. Adv. 9, eadh8502 (2023).

10. Nagamachi, A. et al. Haploinsufficiency of SAMD9L, an endosome fusion facilitator, causes myeloid malignancies in mice mimicking human diseases with monosomy 7. Cancer Cell 24, 305–317 (2013).

11. Narumi, S. et al. SAMD9 mutations cause a novel multisystem disorder, MIRAGE syndrome, and are associated with loss of chromosome 7. Nat. Genet. 48, 792–797 (2016).

12. Hou, G. et al. SAMD9 senses cytosolic double-stranded nucleic acids in epithelial and mesenchymal cells to induce antiviral immunity. Nat. Commun. 16, 3756 (2025).

13. de Jesus, A. A. et al. Distinct interferon signatures and cytokine patterns define additional systemic autoinflammatory diseases. J. Clin. Invest. 130, 1669–1682 (2020).

14. Gorcenco, S. et al. Ataxia-pancytopenia syndrome with SAMD9L mutations. Neurol. Genet. 3, e183 (2017).

15. Topaz, O. et al. A Deleterious Mutation in SAMD9 Causes Normophosphatemic Familial Tumoral Calcinosis. Am. J. Hum. Genet. 79, 759–764 (2006).

16. Wein, T. & Sorek, R. Bacterial origins of human cell-autonomous innate immune mechanisms. Nat. Rev. Immunol. 22, 629–638 (2022).

17. Rousset, F. Innate immunity: the bacterial connection. Trends Immunol. 44, 945–953 (2023).

18. Bernheim, A., Cury, J. & Poirier, E. Z. The immune modules conserved across the tree of life: Towards a definition of ancestral immunity. PLoS Biol. 22, e3002717 (2024).

19. Ledvina, H. E. & Whiteley, A. T. Conservation and similarity of bacterial and eukaryotic innate immunity. Nat. Rev. Microbiol. 22, 420–434 (2024).

20. Mekhedov, S. L., Makarova, K. S. & Koonin, E. V. The complex domain architecture of SAMD9 family proteins, predicted STAND-like NTPases, suggests new links to inflammation and apoptosis. Biol. Direct 12, 13 (2017).

21. Peng, S. et al. Structure and function of an effector domain in antiviral factors and tumor suppressors SAMD9 and SAMD9L. Proc. Natl. Acad. Sci. 119, e2116550119 (2022).

22. Lemos de Matos, A., Liu, J., McFadden, G. & Esteves, P. J. Evolution and divergence of the mammalian SAMD9/SAMD9L gene family. BMC Evol. Biol. 13, 121 (2013).

23. Peeters, M. et al. Risk to Human Health from a Plethora of Simian Immunodeficiency Viruses in Primate Bushmeat. Emerg. Infect. Dis. 8, 451–457 (2002).

24. Chahroudi, A., Bosinger, S. E., Vanderford, T. H., Paiardini, M. & Silvestri, G. Natural SIV Hosts: Showing AIDS the Door. Science 335, 1188–1193 (2012).

25. Li, Y. et al. Eastern chimpanzees, but not bonobos, represent a simian immunodeficiency virus reservoir. J. Virol. 86, 10776–10791 (2012).

26. Van Dooren, S. et al. Lack of evidence for infection with simian immunodeficiency virus in bonobos. AIDS Res. Hum. Retroviruses 18, 213–216 (2002).

27. Keele, B. F. et al. Chimpanzee reservoirs of pandemic and nonpandemic HIV-1. Science 313, 523–526 (2006).

28. Neil, S. J. D., Zang, T. & Bieniasz, P. D. Tetherin inhibits retrovirus release and is antagonized by HIV-1 Vpu. Nature 451, 425–430 (2008).

29. Sawyer, S. L., Wu, L. I., Emerman, M. & Malik, H. S. Positive selection of primate TRIM5α identifies a critical species-specific retroviral restriction domain. Proc. Natl. Acad. Sci. 102, 2832–2837 (2005).

30. Duggal, N. K. & Emerman, M. Evolutionary conflicts between viruses and restriction factors shape immunity. Nat. Rev. Immunol. 12, 687–695 (2012).

31. Etienne, L. et al. The Role of the Antiviral APOBEC3 Gene Family in Protecting Chimpanzees against Lentiviruses from Monkeys. PLoS Pathog. 11, e1005149 (2015).

32. Compton, A. A., Malik, H. S. & Emerman, M. Host gene evolution traces the evolutionary history of ancient primate lentiviruses. Philos. Trans. R. Soc. B Biol. Sci. 368, 20120496 (2013).

33. van Kempen, M. et al. Fast and accurate protein structure search with Foldseek. Nat. Biotechnol. 42, 243– 246 (2024).

34. Tesson, F. et al. A Comprehensive Resource for Exploring Antiphage Defense: DefenseFinder Webservice, Wiki and Databases. 2024.01.25.577194 Preprint at 10.1101/2024.01.25.577194 (2024).

35. Wang, Y., Tian, Y., Yang, X., Yu, F. & Zheng, J. Filamentation activates bacterial Avs5 antiviral protein. Nat. Commun. 16, 2408 (2025).

36. Béchon, N. et al. Diversification of molecular pattern recognition in bacterial NLR-like proteins. Nat. Commun. 15, 9860 (2024).

37. Gao, L. A. et al. Prokaryotic innate immunity through pattern recognition of conserved viral proteins. Science 377, eabm4096 (2022).

38. Gao, L. et al. Diverse enzymatic activities mediate antiviral immunity in prokaryotes. Science 369, 1077–1084 (2020).

39. Rousset, F. et al. A conserved family of immune effectors cleaves cellular ATP upon viral infection. Cell 186, 3619–3631.e13 (2023).

40. Garb, J. et al. Multiple phage resistance systems inhibit infection via SIR2-dependent NAD+ depletion. Nat. Microbiol. 7, 1849–1856 (2022).

41. Schultz, J., Ponting, C. P., Hofmann, K. & Bork, P. SAM as a protein interaction domain involved in developmental regulation. Protein Sci. Publ. Protein Soc. 6, 249–253 (1997).

42. Xu, J. & Zhang, Y. How significant is a protein structure similarity with TM-score = 0.5? Bioinformatics 26, 889–895 (2010).

43. Leipe, D. D., Koonin, E. V. & Aravind, L. STAND, a class of P-loop NTPases including animal and plant regulators of programmed cell death: multiple, complex domain architectures, unusual phyletic patterns, and evolution by horizontal gene transfer. J. Mol. Biol. 343, 1–28 (2004).

44. Loquet, A. & Saupe, S. J. Diversity of Amyloid Motifs in NLR Signaling in Fungi. Biomolecules 7, 38 (2017).

45. Mordier, J., Fraisse, M., Cohen-Tannoudji, M. & Molaro, A. Recurrent Evolutionary Innovations in Rodent and Primate Schlafen Genes. 2024.01.12.575368 Preprint at 10.1101/2024.01.12.575368 (2024).

46. Doron, S. et al. Systematic discovery of anti-phage defense systems in the microbial pan-genome. Science 359, eaar4120 (2018).

47. Valenzuela, C., Saucedo, S. & Llano, M. Schlafen14 Impairs HIV-1 Expression in a Codon Usage-Dependent Manner. Viruses 16, 502 (2024).

48. Kobayashi-Ishihara, M. et al. Schlafen 12 restricts HIV-1 latency reversal by a codon-usage dependent post-transcriptional block in CD4+ T cells. *Commun*. Biol. 6, 1–15 (2023).

49. Metzner, F. J., Huber, E., Hopfner, K.-P. & Lammens, K. Structural and biochemical characterization of human Schlafen 5. Nucleic Acids Res. 50, 1147–1161 (2022).

50. Garvie, C. W. et al. Structure of PDE3A-SLFN12 complex reveals requirements for activation of SLFN12 RNase. Nat. Commun. 12, (2021).

51. Yang, J.-Y. et al. Structure of Schlafen13 reveals a new class of tRNA/rRNA-targeting RNase engaged in translational control. Nat. Commun. 9, 1165 (2018).

52. Li, M. et al. DNA damage-induced cell death relies on SLFN11-dependent cleavage of distinct type II tRNAs. Nat. Struct. Mol. Biol. 25, 1047–1058 (2018).

53. Picard, L. et al. DGINN, an automated and highly-flexible pipeline for the detection of genetic innovations on protein-coding genes. Nucleic Acids Res. 48, e103 (2020).

54. Wellenreuther, M., Mérot, C., Berdan, E. & Bernatchez, L. Going beyond SNPs: The role of structural genomic variants in adaptive evolution and species diversification. Mol. Ecol. 28, 1203–1209 (2019).

55. Radke, D. W. & Lee, C. Adaptive potential of genomic structural variation in human and mammalian evolution. Brief. Funct. Genomics 14, 358–368 (2015).

56. Daugherty, M. D. & Malik, H. S. Rules of Engagement: Molecular Insights from Host-Virus Arms Races. Annu. Rev. Genet. 46, 677–700 (2012).

57. Smith, M. D. et al. Less Is More: An Adaptive Branch-Site Random Effects Model for Efficient Detection of Episodic Diversifying Selection. Mol. Biol. Evol. 32, 1342–1353 (2015).

58. Prüfer, K. et al. The bonobo genome compared with the chimpanzee and human genomes. Nature 486, 527–531 (2012).

59. Mao, Y. et al. A high-quality bonobo genome refines the analysis of hominid evolution. Nature 594, 77–81 (2021).

60. Prado-Martinez, J. et al. Great ape genetic diversity and population history. Nature 499, 471–475 (2013).

61. Allenspach, E. J. et al. Germline SAMD9L truncation variants trigger global translational repression. J. Exp. Med. 218, e20201195 (2021).

62. Thomas, M. E. et al. Pediatric MDS and bone marrow failure-associated germline mutations in SAMD9 and SAMD9L impair multiple pathways in primary hematopoietic cells. Leukemia 35, 3232–3244 (2021).

63. Locatelli, S. et al. Why Are Nigeria-Cameroon Chimpanzees (Pan troglodytes ellioti) Free of SIVcpz Infection? PLOS ONE 11, e0160788 (2016).

64. Gao, F. et al. Origin of HIV-1 in the chimpanzee Pan troglodytes troglodytes. Nature 397, 436–441 (1999).

65. Barbian, H. J. et al. Neutralization properties of simian immunodeficiency viruses infecting chimpanzees and gorillas. mBio 6, e00296–15 (2015).

66. Burroughs, A. M. & Aravind, L. Identification of Uncharacterized Components of Prokaryotic Immune Systems and Their Diverse Eukaryotic Reformulations. J. Bacteriol. 202, 10.1128/jb.00365-20 (2020).

67. Sawyer, S. L., Emerman, M. & Malik, H. S. Ancient Adaptive Evolution of the Primate Antiviral DNA-Editing Enzyme APOBEC3G. PLoS Biol. 2, e275 (2004).

68. Münk, C., Willemsen, A. & Bravo, I. G. An ancient history of gene duplications, fusions and losses in the evolution of APOBEC3 mutators in mammals. BMC Evol. Biol. 12, 71 (2012).

69. Ito, J., Gifford, R. J. & Sato, K. Retroviruses drive the rapid evolution of mammalian APOBEC3 genes. Proc. Natl. Acad. Sci. U. S. A. 117, 610–618 (2020).

70. Kim, C. A. & Bowie, J. U. SAM domains: uniform structure, diversity of function. Trends Biochem. Sci. 28, 625–628 (2003).

71. Knight, M. J., Leettola, C., Gingery, M., Li, H. & Bowie, J. U. A human sterile alpha motif domain polymerizome. Protein Sci. 20, 1697–1706 (2011).

72. Johnson, B. et al. Human IFIT3 Modulates IFIT1 RNA Binding Specificity and Protein Stability. Immunity 48, 487–499.e5 (2018).

73. Fleith, R. C. et al. IFIT3 and IFIT2/3 promote IFIT1-mediated translation inhibition by enhancing binding to non-self RNA. Nucleic Acids Res. 46, 5269–5285 (2018).

74. Sahoo, S. S. et al. Clinical evolution, genetic landscape and trajectories of clonal hematopoiesis in SAMD9/SAMD9L syndromes. Nat. Med. 27, 1806–1817 (2021).

75. Keele, B. F. et al. Increased mortality and AIDS-like immunopathology in wild chimpanzees infected with SIVcpz. Nature 460, 515–519 (2009).

76. Etienne, L. et al. Characterization of a new simian immunodeficiency virus strain in a naturally infected Pan troglodytes troglodytes chimpanzee with AIDS related symptoms. Retrovirology 8, 4 (2011).

77. Greenwood, E. J. D. et al. Simian Immunodeficiency Virus Infection of Chimpanzees (Pan troglodytes) Shares Features of Both Pathogenic and Non-pathogenic Lentiviral Infections. PLOS Pathog. 11, e1005146 (2015).

78. Pawar, H., Ostridge, H. J., Schmidt, J. M. & Andrés, A. M. Genetic adaptations to SIV across chimpanzee populations. PLoS Genet. 18, e1010337 (2022).

79. Camacho, C. et al. BLAST+: architecture and applications. BMC Bioinformatics 10, 421 (2009).

80. Altschul, S. F., Gish, W., Miller, W., Myers, E. W. & Lipman, D. J. Basic local alignment search tool. J. Mol. Biol. 215, 403–410 (1990).

81. Katoh, K. & Standley, D. M. MAFFT multiple sequence alignment software version 7: improvements in performance and usability. Mol. Biol. Evol. 30, 772–780 (2013).

82. Löytynoja, A. Phylogeny-aware alignment with PRANK. Methods Mol. Biol. Clifton NJ 1079, 155–170 (2014).

83. Guindon, S. et al. New Algorithms and Methods to Estimate Maximum-Likelihood Phylogenies: Assessing the Performance of PhyML 3.0. Syst. Biol. 59, 307–321 (2010).

84. Edgar, R. C. MUSCLE: multiple sequence alignment with high accuracy and high throughput. Nucleic Acids Res. 32, 1792–1797 (2004).

85. Trifinopoulos, J., Nguyen, L.-T., von Haeseler, A. & Minh, B. Q. W-IQ-TREE: a fast online phylogenetic tool for maximum likelihood analysis. Nucleic Acids Res. 44, W232–W235 (2016).

86. Bergström, A. et al. Insights into human genetic variation and population history from 929 diverse genomes. Science 367, eaay5012 (2020).

87. Li, H. Aligning sequence reads, clone sequences and assembly contigs with BWA-MEM. Preprint at 10.48550/arXiv.1303.3997 (2013).

88. Garrison, E. & Marth, G. Haplotype-based variant detection from short-read sequencing. Preprint at 10.48550/arXiv.1207.3907 (2012).

89. Jumper, J. et al. Highly accurate protein structure prediction with AlphaFold. Nature 596, 583–589 (2021).

90. Varadi, M. et al. AlphaFold Protein Structure Database: massively expanding the structural coverage of protein-sequence space with high-accuracy models. Nucleic Acids Res. 50, D439–D444 (2022).

91. Berman, H. M. et al. The Protein Data Bank. Nucleic Acids Res. 28, 235–242 (2000).

92. Gilchrist, C. L. M., Mirdita, M. & Steinegger, M. Multiple Protein Structure Alignment at Scale with FoldMason. 2024.08.01.606130 Preprint at 10.1101/2024.08.01.606130 (2024).

93. Minh, B. Q. et al. IQ-TREE 2: New Models and Efficient Methods for Phylogenetic Inference in the Genomic Era. Mol. Biol. Evol. 37, 1530–1534 (2020).

94. Yu, G., Smith, D. K., Zhu, H., Guan, Y. & Lam, T. T.-Y. ggtree: an r package for visualization and annotation of phylogenetic trees with their covariates and other associated data. Methods Ecol. Evol. 8, 28–36 (2017).

95. Letunic, I. & Bork, P. Interactive Tree of Life (iTOL) v6: recent updates to the phylogenetic tree display and annotation tool. Nucleic Acids Res. 52, W78–W82 (2024).

96. Paysan-Lafosse, T. et al. The Pfam protein families database: embracing AI/ML. Nucleic Acids Res. 53, D523–D534 (2025).

97. Steinegger, M. & Söding, J. MMseqs2 enables sensitive protein sequence searching for the analysis of massive data sets. Nat. Biotechnol. 35, 1026–1028 (2017).

98. Richter, D. J. et al. EukProt: A database of genome-scale predicted proteins across the diversity of eukaryotes. Peer Community J. 2, (2022).

99. Eddy, S. R. Accelerated Profile HMM Searches. PLoS Comput. Biol. 7, e1002195 (2011).

100. Sievers, F. et al. Fast, scalable generation of high-quality protein multiple sequence alignments using Clustal Omega. Mol. Syst. Biol. 7, 539 (2011).

101. Steenwyk, J. L., Iii, T. J. B., Li, Y., Shen, X.-X. & Rokas, A. ClipKIT: A multiple sequence alignment trimming software for accurate phylogenomic inference. PLOS Biol. 18, e3001007 (2020).

102. Lee, T. S. et al. BglBrick vectors and datasheets: A synthetic biology platform for gene expression. J. Biol. Eng. 5, 12 (2011).

103. Xia, Y. et al. T5 exonuclease-dependent assembly offers a low-cost method for efficient cloning and site-directed mutagenesis. Nucleic Acids Res. 47, e15 (2019).

104. Doyle, T. et al. The interferon-inducible isoform of NCOA7 inhibits endosome-mediated viral entry. Nat. Microbiol. 3, 1369–1376 (2018).

105. Robert, X. & Gouet, P. Deciphering key features in protein structures with the new ENDscript server. Nucleic Acids Res. 42, W320–W324 (2014).

106. Okonechnikov, K., Golosova, O., Fursov, M., & UGENE team. Unipro UGENE: a unified bioinformatics toolkit. Bioinforma. Oxf. Engl. 28, 1166–1167 (2012).

