## Supplementary figures and images for "Ancient convergence with prokaryote defense and recent adaptations to lentiviruses in primates characterize the ancestral immune factors SAMD9s"

### Datase S2

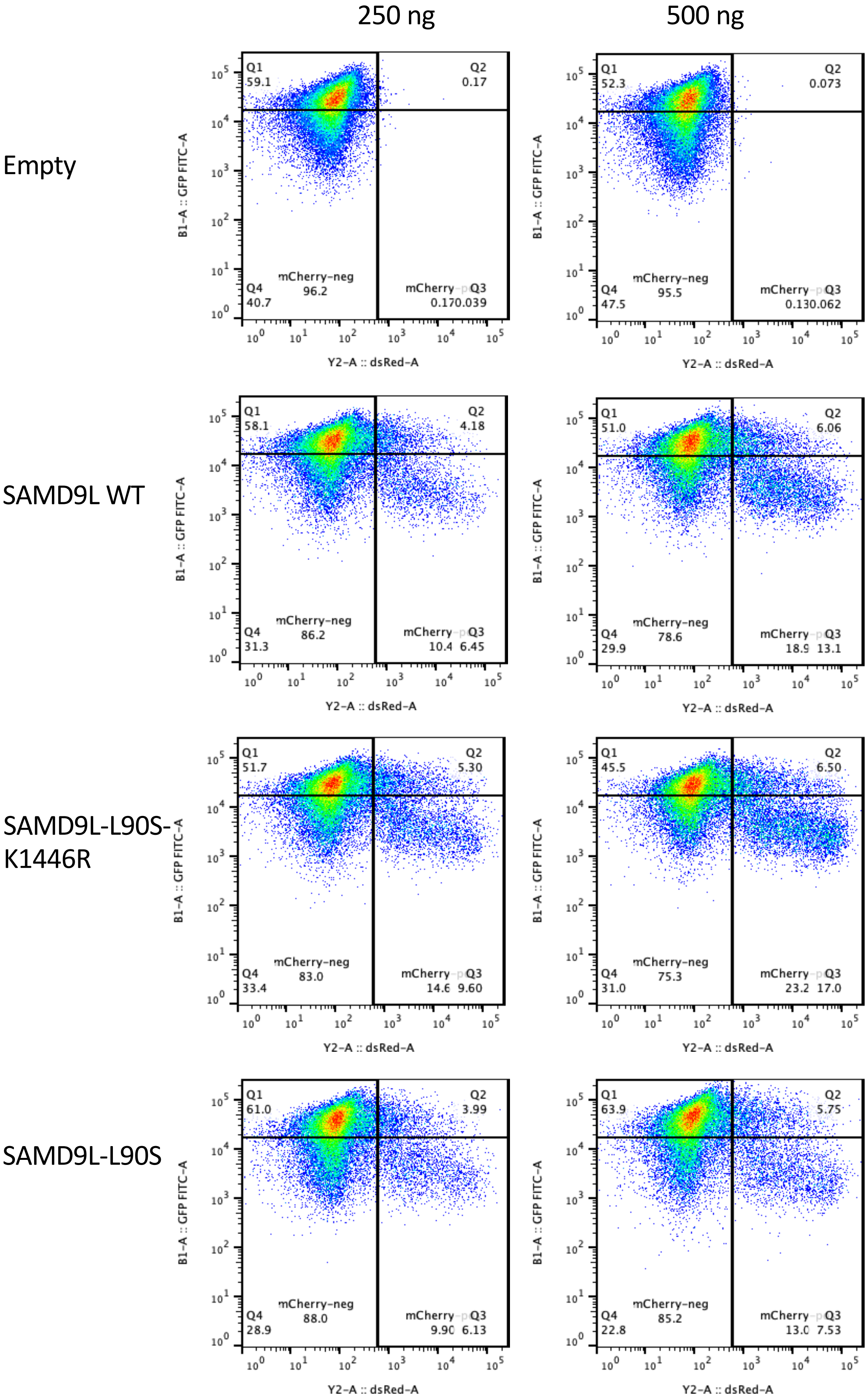

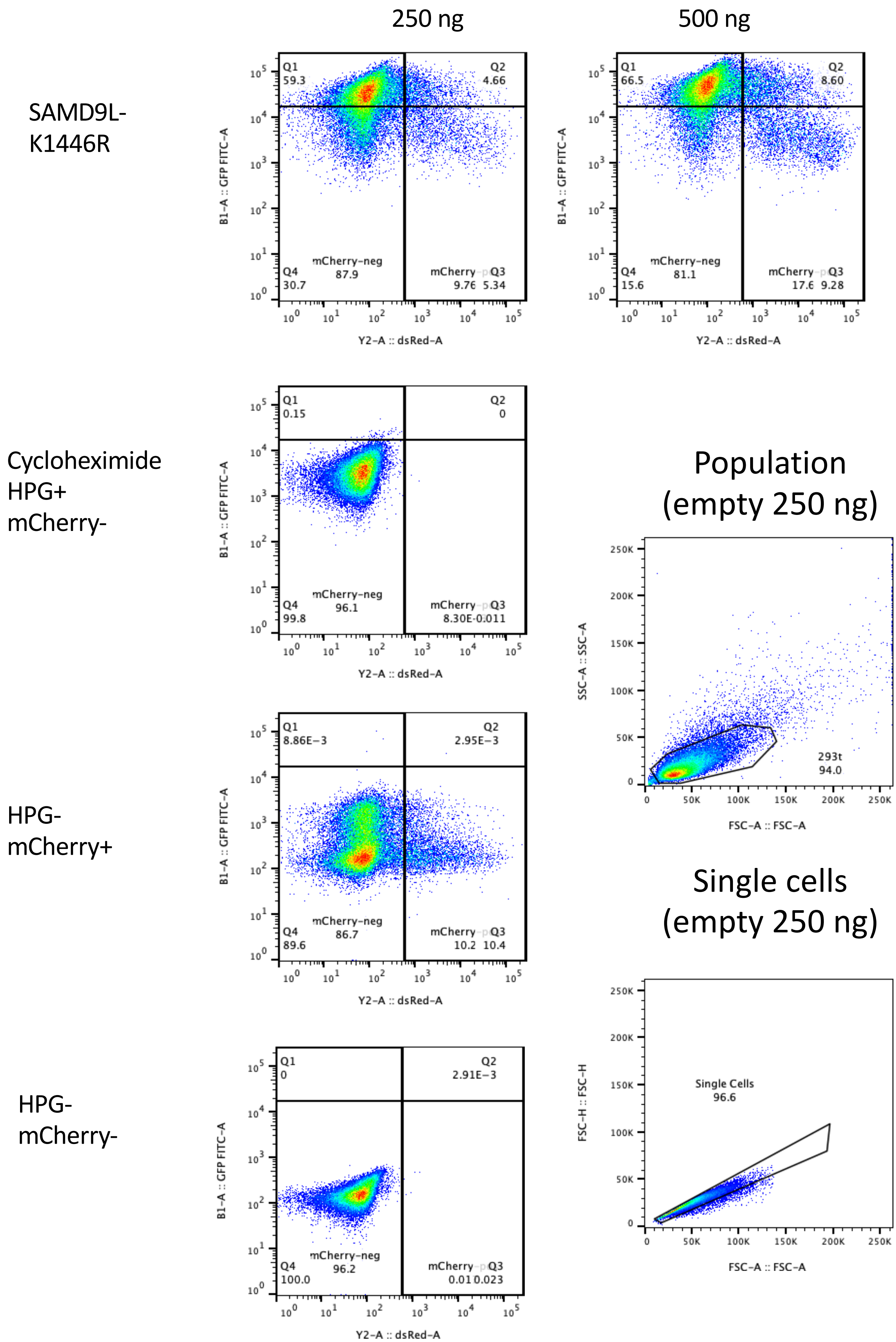
